# Behavioral responses buffer seasonal variation more strongly than endocrine and chemical responses in poison frogs

**DOI:** 10.64898/2026.03.14.711838

**Authors:** Shirley Jennifer Serrano-Rojas, Andrius Pašukonis, Mabel Gonzalez, Camilo Rodriguez, Rodrigo F. Calvo Usto, Andrea Carazas, Cristel Sandoval García, Jean Pier Zolorzano, Luisa F. Arcila-Pérez, Sergio Boluarte-Salinas, Esaú Baldarrago, Alfredo Sosa-Salazar, Lauren A. O’Connell

## Abstract

Seasonal rainfall shapes biological responses in tropical ecosystems, yet how tropical organisms integrate behavioral and physiological responses to cope with seasonality remains poorly understood. We assessed how four poison frog species with contrasting reproductive strategies respond to dry and wet season environmental conditions. We quantified spatial behavior, microhabitat use, hormone release rates, and chemical defenses in two seasonal breeders (*Allobates femoralis* and *Ameerega trivittata*) and two year-round breeders (*Ameerega macero* and *Ameerega shihuemoy*). Seasonal breeders exhibited pronounced sex-specific shifts in space use, where males expanded their space use during the wet season, likely to track reproductive opportunities, while *A. femoralis* females increased their spatial use during the dry season, likely responding to foraging demands when prey resources are sparse. Year-round breeders maintained similar space use across seasons, likely reflecting their ability to access key resources within the same space to reproduce year-round. Microhabitat use was flexible, as seasonal breeders shifted toward humid refugia during the dry season and reproduction-associated microhabitats during the wet season, whereas year-round breeders selected microhabitats that facilitate continuous reproduction across seasons. Despite these behavioral responses, water-borne corticosterone and testosterone, as well as chemical defenses, showed no consistent seasonal variation, suggesting that behavioral responses may be partially decoupled from shifts in endocrine and chemical defenses. These results support a role for behavioral buffering in mediating responses to seasonal environmental variation. However, given the increasing unpredictability in rainfall timing and intensity as a result of climate change, how these coping strategies will function in the long term is uncertain.

## Introduction

Climate change drives biodiversity loss (Pfenning-Butterworth et al. 2024), partially by altering the timing, intensity, and predictability of environmental factors like temperature and rainfall (Webb et al. 2005). For some species, seasonal rainfall strongly shapes resource availability, reproductive timing, and environmental stress (Feng et al. 2013). Tropical ecosystems exhibit pronounced rainfall seasonality, with distinct dry and wet seasons that shape biological and ecological processes (Brown and Shine 2006; Mandl et al. 2018). Thus, seasonal habitats provide a natural context to understand how organisms coordinate behavioral and physiological responses to climatic variability (Varpe 2017; Gotthard et al. 2025). Understanding these mechanisms is critical for predicting resilience or vulnerability to climate change.

Contrasting life-history traits provide a comparative framework for understanding how species respond to seasonal variability (Hjernquist et al. 2012; Hall and MacPherson 2025). Reproductive timing, for example, ranges from reproducing during specific periods of favorable conditions in a season to reproducing continuously throughout the year (Ndithia et al. 2017; Heldstab et al. 2021). Seasonal breeders minimize exposure to unfavorable conditions by limiting their reproduction to periods of high resource availability (Hall et al. 2018; Sun et al. 2020; Soanes et al. 2021). In contrast, year-round breeders exhibit behaviors that may reduce exposure to seasonal constraints while maintaining reproductive activity across fluctuating environmental conditions (Prado et al. 2005; Heinermann et al. 2015). Thus far, relatively few tropical studies have investigated seasonal responses through a framework that integrates behavior, spatial ecology, and physiology across species with contrasting life-history strategies (Lopez et al. 2023). Behavioral responses to seasonal variability may also occur independently of broad endocrine shifts, highlighting the importance of assessing both behavioral and physiological responses simultaneously. Glucocorticoids and gonadal steroids are often used to assess physiological responses to environmental variation because of their roles on energy allocation, reproduction, and behavior (Crespi et al. 2013; Husak et al. 2021). In amphibians, corticosterone is one of the main glucocorticoids involved in metabolic regulation and responses to environmental change, whereas testosterone influences reproductive behaviors, spatial activity, and aggression (Wingfield et al. 1998; Moore and Jessop 2003; Denver 2009a; Eikenaar et al. 2012; Husak et al. 2021). Assessing seasonal variation in these hormones alongside behavioral responses can therefore reveal whether seasonal environmental change is accompanied by endocrine adjustments. Filling this critical gap is important in the context of global climate change because climate projections for tropical regions consistently predict alterations in the timing, intensity, and predictability of rainfall (Chadwick et al. 2016; Pendergrass et al. 2017; Song et al. 2023).

Across frog species, seasonal rainfall shapes life-history traits (Watling and Donnelly 2002; Ficetola and Maiorano 2016), alters microhabitat selection, movement, and diet (Born et al. 2010), and drives plasticity in parental care (Schulte and Lötters 2013). Poison frogs (Dendrobatidae and Aromobatidae) offer contrasting life-history traits for assessing seasonal responses. Some species breed seasonally and rely on temporary ponds or phytotelmata for tadpole development, while others breed year-round using similar habitats during the wet season and pools along stream margins during the dry season (Wells 2010). Despite these differences in reproduction timing, poison frogs broadly share parental care behaviors and rely on fine-scale spatial behavior and specific microhabitats for reproduction and thermal refuges (Brown et al. 2008). Poison frogs also sequester alkaloids from arthropod prey as chemical defenses (Saporito et al. 2003; Daly et al. 2005), and the composition of these defenses can vary across space and time, reflecting changes in prey communities across seasons (Saporito et al. 2006). Seasonal changes in rainfall may influence chemical defenses by altering prey availability and foraging opportunities. Thus, assessing chemical defenses can provide further insight into how seasonal environmental variation affects poison frogs via trophic interactions and into the potential consequences of predicted rainfall changes in the tropics.

In this study, we investigated how seasonal rainfall affects poison frogs with contrasting reproductive timing. We focused on two seasonal breeders (*Allobates femoralis* and *Ameerega trivittata*), which reproduce mostly during the wet season, and two year-round breeders (*Ameerega macero* and *Ameerega shihuemoy*), which reproduce throughout the year. Here, we define behavioral buffering as behavioral adjustments that reduce exposure to unfavorable environmental conditions, including shifts in space use area, or microhabitat preferences across seasons. We hypothesize that seasonal breeders will show stronger behavioral, physiological, and chemical defense responses to rainfall shifts than year-round breeders. Specifically, we predict that seasonal breeders will reduce space use and seek refuge in humid microhabitats during the dry season. Seasonal environmental changes will be accompanied by shifts in corticosterone and testosterone release rates, thus reflecting physiological responses to changing environmental and reproductive conditions. These environmental changes, together with reduced space use, may also alter access to alkaloid-rich prey and lead to seasonal shifts in chemical defenses. In contrast, year-round breeders are expected to exhibit behavioral buffering, maintaining similar movement patterns, space use, hormone release rates, and chemical defenses across seasons despite variation in rainfall.

## Materials and methods

### Study site and species

We conducted this study in two sites within the Madre de Dios Region, southeastern Peru (Fig. 1A). Across both sites, seasonal variation is primarily driven by rainfall rather than temperature, reflecting the strong precipitation seasonality typical of tropical forest systems.

**Fig. 1.**
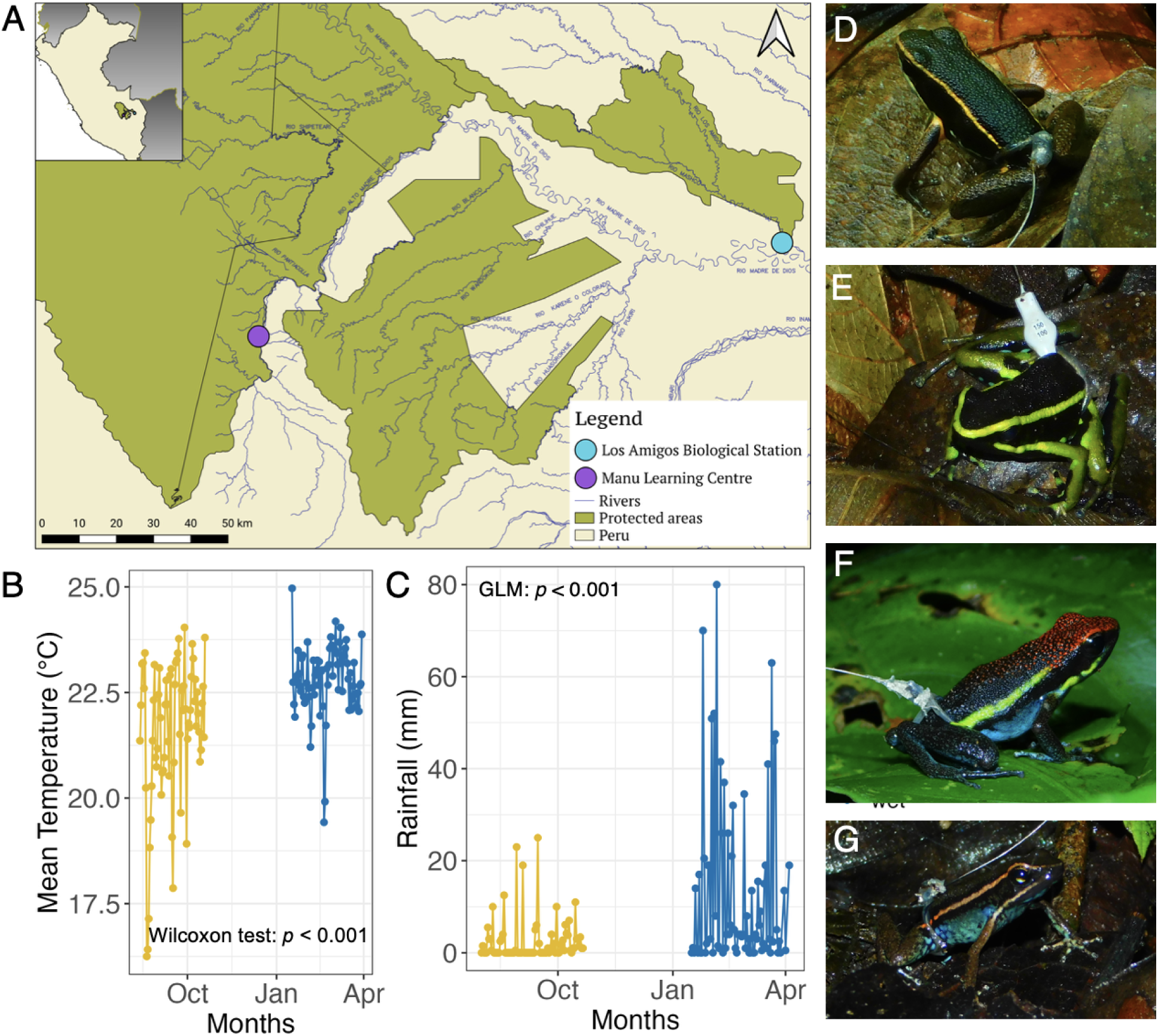
Study sites, environmental conditions, and focal species. (A) Map of southeastern Peru showing the two study sites: the Manu Learning Centre (purple), where *Allobates femoralis*, *Ameerega macero*, and *Ameerega shihuemoy* were studied, and the Los Amigos Biological Station (blue), where *Ameerega trivittata* was studied. (B) Mean temperature and (C) daily precipitation (mm/day) during the dry (yellow) and wet (blue) seasons at the Manu Learning Centre. (D–G) Field photographs of tagged individuals of the focal species: (D) *A. femoralis* (brilliant-thighed poison frog), (E) *A. trivittata* (three-striped poison frog), (F) *A. macero* (Manu poison frog), and (G) *A. shihuemoy* (Amarakaeri poison frog). Comparable environmental data for Los Amigos Biological Station are provided in Supplementary Fig. S1.

Three species (*Allobates femoralis*, *Ameerega macero,* and *Ameerega shihuemoy*) were studied in a regenerating forest within the buffer zone of the Manu Biosphere Reserve (The Manu Learning Centre; 450 – 700 m asl). Based on weather data collected on site during the sampling periods (dry season: August–early October 2022; wet season: January–March 2023), mean daily temperature differed between seasons (Wilcoxon rank-sum: W=1288, p<0.001; Fig. 1B), with the wet season slightly warmer (22.82 °C) than the dry season (22.04 °C). Rainfall probability and intensity were also higher in the wet season (3.75× and ∼3.3× greater, respectively; p < 0.001), and daily rainfall patterns showed pronounced seasonal differences (Fig. 1C).

*Ameerega trivittata* was studied at Los Amigos Biological Station (300 m asl), 142 km from Manu. Based on weather data collected on site during the sampling periods (dry season: November 2022; wet season: January–February 2024), this floodplain forest showed no seasonal difference in mean daily temperature (Welch t-test: t=0.10, p=0.92; dry = 25.14 °C, wet = 25.11 °C), but rainfall probability was 6.72× higher in the wet season ( *p* < 0.001; Supplementary Methods S1 and Supplementary Fig. S1).

All four species share similar mating and parental behaviors (Rodríguez and Myers 1993; Roithmair 1994; Kaefer et al. 2012; Serrano-Rojas et al. 2017; Fig. 1D–G). Based on breeding duration, we classified *A. femoralis* and *A. trivittata* as seasonal breeders and *A. macero* and *A. shihuemoy* as year-round breeders (see Supplementary Methods S2 and Fig. S2). The four focal species belong to the superfamily Dendrobatoidea but differ in their phylogenetic relatedness. Three species (*A. macero*, *A. shihuemoy,* and *A. trivittata*) are members of the family Dendrobatidae and the genus Ameerega (Guillory et al. 2020). In contrast, *A. femoralis* belongs to the family Aromobatidae, which diverged from Dendrobatidae approximately 40 million years ago (Santos et al. 2009). While we do not account for phylogenetic relatedness in our analyses, this context provides a comparative framework for understanding species differences.

### Telemetry, space-use quantification, and microhabitat use

We selected telemetry methods based on body size. Smaller species (*A. femoralis, A. macero, and A. shihuemoy*) were tracked using harmonic direction finding (HDF) with passive transponders, whereas the larger *A. trivittata* was tracked using radio-telemetry with very high frequency (VHF) transmitters following Pašukonis et al. (2014, 2018, 2022). See Supplementary Methods S3 for details and a step-by-step protocol available on protocols.io (Serrano-Rojas et al. 2026).

We tracked 212 frogs across species. Individuals were located 5–7 times daily, recording position and perching substrate. When direct observation was not possible due to dense vegetation, we estimated locations within ∼1 m. The tracking area was mapped using compasses and laser distance meters to establish reference points (Ringler et al. 2016). Distances and bearings were recorded using the Epicollect5 app (Centre for Genomic Pathogen Surveillance 2025) and converted to UTM (Universal Transverse Mercator) coordinates in R and QGIS.

We excluded all locations associated with tadpole transport because these movements are highly directional, faster, and cover longer distances than routine movements, which would inflate space-use estimates. Additionally, we excluded tracking days with fewer than four locations and individuals tracked for fewer than five full days, because sampling rate and tracking duration influence spatial parameters.

We defined microhabitat as the structural substrate and spatial position occupied at each relocation. We categorized substrates into 13 categories. Exposed substrates (e.g., leaf litter, leaves, branches, logs, palm roots, rocks) were frequently used for calling, perching, and moving. Potential clutch sites were those previously observed to be used for egg deposition, such as palm bracts and rolled leaves that form sheltered cavities. Refuge or concealed microhabitats (e.g., root cavities, tree fall areas, rock or soil cavities, and spaces under logs or leaf litter) were primarily used during the hottest periods of the day, although recent treefalls also served as calling sites.

### Water-borne hormones

We measured water-borne corticosterone and testosterone release rates in individuals tracked for ≥5 days following an indirect non-invasive water-borne hormone sampling method originally developed for amphibians by Gabor et al. (2013) and subsequently adapted for other frog species, including poison frogs (Baugh et al. 2018; Rodríguez et al. 2022). Briefly, we captured frogs between 15:00 and 18:00 h and placed them individually in glass containers containing 40 mL of distilled water for 1 h, allowing hormones released through the skin and urine to accumulate in the water. Water samples were then filtered through C18 cartridges and hormones were eluted with ethanol for subsequent laboratory analysis. See Supplementary Methods S4 for extraction and processing details and Table S1 for sample sizes.

We quantified corticosterone and testosterone using commercial enzyme immunoassay kits (corticosterone: ADI-900-097; testosterone: ADI-900-065, Enzo Life Sciences, Farmingdale, NY, USA). We converted the hormone concentrations to total hormone released and expressed as water-borne hormone release rates (pg/h; see Supplementary Methods S4). To minimize batch effects, we randomized the order of sample assays, ensuring each batch included individuals from all species and seasons within the same geographical location. Due to differences in sampling timing, *A. trivittata* samples were processed separately. Since our analysis focuses on within-species seasonal comparisons, this approach does not affect our main inferences. Samples with a coefficient of variation (CV) > 20% were excluded (corticosterone: 38/145; testosterone: 29/145). Intra-assay CVs were 8.27% and 7.38%, and inter-assay CVs were 6.77% and 10.17%, respectively.

### Alkaloid collection, extraction, and annotation

We used non-lethal metabolite profiling of skin secretions following Gonzalez (2021) to extract and quantify alkaloids in frogs tracked for ≥5 days (see Supplementary Methods S5 for details). Alkaloids were analyzed using gas chromatography/mass spectrometry (GC/MS) following Gonzalez et al. (2021) and annotated using the Global Natural Products Social Molecular Networking (GNPS) pipeline (Supplementary Methods S6; Supplementary Table S2).

## Statistical analyses

All statistics were conducted in R (version 2025.09.0). While statistical methods are briefly described here, a complete description can be found in the Supplementary Materials. Code is available on the GitHub repository (https://github.com/laurenoconnelllab/Behavioral-responses-buffer-seasonal-variation-m ore-strongly-than-endocrine-and-chemical-responses).

### Space use and daily movement

We estimated individual space use (m²) as the 95% utilization distribution (UD) derived from kernel density estimation (KDE), using the *kernelUD()* function from the *adehabitatHR* R package (Calenge 2006) and a conservative plug-in bandwidth selection method with the *hpi()* function from the *ks* R package (Chacón and Duong 2018). Most individuals were sampled in only one season, although fourteen individuals were recaptured across seasons. Therefore, seasonal comparisons primarily represent population-level changes in space use rather than within-individual seasonal changes. For each species, we tested the effects of season and sex, and their interaction, on space use area using linear models, followed by post hoc tests (Supplementary Methods S7). Models included snout–vent length (SVL) and tracking duration as covariates to account for body-size differences that may influence movement capacity and variation in sampling effort among individuals, respectively. To explore seasonal shifts in location, we quantified seasonal space use overlap and centroid shifts for recaptured individuals (see detailed methods in Supplementary Methods S8).

As tadpole transport events were recorded unevenly across individuals, including them in space use area estimates would have introduced sampling bias and reduced comparability among individuals. However, we provide complementary analysis including tadpole transport in Supplementary Methods S9.

Additionally, we estimated individual daily movement and assessed how rainfall and temperature influence daily movement using linear mixed-effect models for each species (Supplementary Methods S10).

### Microhabitat use

To assess seasonal differences in microhabitat use, we fitted generalized linear mixed-effects models (GLMMs) with negative binomial error distribution (nbinom2) using the *glmmTMB()* function from the *glmmTMB* R package (Brooks et al. 2026) for each species. Full model details are in Supplementary Methods S11.

### Water-borne hormone release rates

We assessed seasonal differences in hormonal release rates using species-specific models. Testosterone and corticosterone release rates (pg/h) were log-transformed and analyzed with linear models, except for *A. macero* (testosterone) and *A. trivittata* (corticosterone), which required Gamma GLMs with log links. For species with data for both sexes, we assessed whether seasonal effects depended on sex while accounting for body size (log(hormone) ∼ season*sex + SVL); otherwise, we only included season and SVL. Significant effects were evaluated using estimated marginal means with FDR-corrected comparisons. Results were back-transformed for visualization. Additionally, we assessed the influence of testosterone release rates, sex, and their interaction on space use area by using linear models for each species.

### Alkaloids

We normalized alkaloid peak intensities using nicotine internal standard (IS) (Grant et al. 2012; Caty et al. 2025). We then grouped alkaloids into structural families based on revised annotations (Supplementary Table S2). We quantified alkaloid family-level abundance (ng), alkaloid richness (number of detected alkaloids), and total alkaloid abundance for each individual frog. We also quantified detection frequency per alkaloid family within species and seasons. To evaluate seasonal differences in overall alkaloid composition, we constructed a Bray–Curtis dissimilarity matrix using alkaloid family-level abundances and performed species-specific permutational multivariate analyses of variance (PERMANOVA) using the *adonis2()* function in the *vegan* R package (Oksanen et al. 2025) with 999 permutations. Homogeneity of multivariate dispersion was evaluated using *betadisper()*. To assess seasonal differences in individual alkaloid families within species, we used Wilcoxon rank-sum tests and adjusted p-values for multiple comparisons using the Benjamini–Hochberg procedure. For visualization, we summarized alkaloid abundance using mean abundance and 95% confidence intervals and included individual-level observations for frogs with detectable alkaloids.

## Results

### Seasonal and sex differences in space use area reflect breeding strategies

We evaluated whether season influences space use area in species with contrasting breeding strategies (Fig. 2A). When considering all individuals (i.e., both males and females; Fig. 2A, upper panel), *A. trivittata* and *A. macero* showed slightly larger space use area in the wet season, whereas *A. femoralis* and *A. shihuemoy* showed slightly larger space use area in the dry season. However, statistically significant seasonal differences were only detected in *A. trivittata*. In the seasonal breeder *A. femoralis*, space use area showed sex-specific seasonal patterns (LM: season*sex: F_1,50_=12.99, *p* < 0.001). Tracking duration, included as a covariate in the model, also showed a significant positive effect on space use area (p < 0.05). Females showed larger space use area in the dry season (emmeans estimate = 1.28 ± 0.50, *p_adj_* = 0.014), while males exhibited smaller space use area in the dry season (estimate = –0.84 ± 0.35, *p_adj_* = 0.021). In the seasonal breeder *A. trivittata*, space use area differed between seasons regardless of sex (LM: season: F_1,33_=6.02, *p* < 0.020), with individuals exhibiting smaller space use area in the dry season (estimate = –0.75 ± 0.30, *p_adj_* = 0.017). In the year-round breeder *A. macero*, space use area showed a season*sex interaction (LM: F_1,49_=4.71, *p* = 0.035); however, we detected no significant seasonal shift for either sex. The other year-round breeder, *A. shihuemoy*, showed no seasonal variation in space use area. Full model outputs and post hoc comparisons are provided in Supplementary Tables S3 and S4. Seasonal space use overlap and centroid shifts varied among species and sexes (Supplementary Results S1; Supplementary Tables S5 and S6). However, because the number of recaptures was limited, these results should be interpreted as exploratory only. The inclusion of tadpole transport movements in space use area estimates did not alter the overall conclusions (Supplementary Results S2; Supplementary Table S7).

**Fig. 2.**
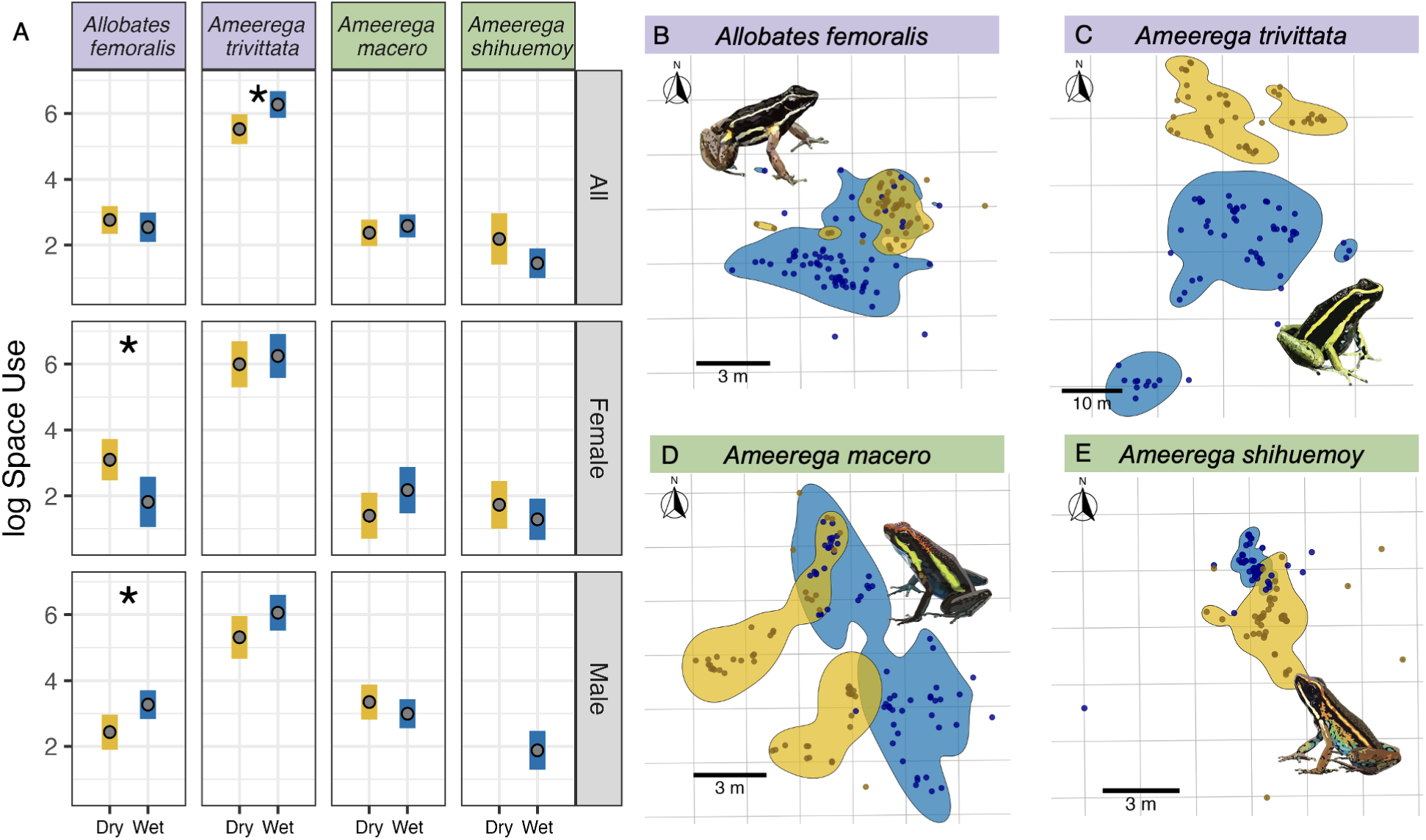
Species-specific seasonal shifts in space use area reflect contrasting breeding strategies in poison frogs. (A) Seasonal differences in space use area between sexes and across species. Points represent estimated marginal means, and bars represent the 95% confidence intervals from the selected linear models. Asterisks indicate significant differences ( * < 0.05). (B–E) Kernel density estimates illustrating the spatial distribution and overlap of seasonal space use in recaptured individual: (B) male *Allobates femoralis,* (C) male *Ameerega trivittata*, (D) male *Ameerega macero,* and (E) female *Ameerega shihuemoy* (no male recaptures were obtained for this species). Yellow indicates the dry season and blue indicates the wet season. Purple panels represent seasonal breeders and green panels represent year-round breeders. Sample sizes for panel A were: *A. femoralis* dry season (12 females, 14 males) and wet season (7 females, 23 males); *A. trivittata* dry (7 females, 9 males) and wet (8 females, 12 males); *A. macero* dry (8 females, 13 males) and wet (11 females, 22 males); and *A. shihuemoy* dry (6 females, 1 male) and wet (8 females, 9 males).

Environmental variables also influenced daily movement differently among species: rainfall was associated with movement in *A. femoralis*, temperature influenced movement in *A. trivittata* and *A. macero*, and no effects were detected for *A. shihuemoy* (Supplementary Table S8 and Fig. S3).

### Species-specific seasonal microhabitat shifts

In the seasonal breeder *A. femoralis,* seasonal changes in microhabitat differed between sexes for specific substrate types (GLMM: season*sex*microhabitat: *χ2*(8) = 22.37, *p* = 0.004; Supplementary Table S9). Post hoc contrasts (Supplementary Table S10) revealed that females increased use of exposed structures (e.g., logs) during the wet season (dry/wet ratio = 0.56, *p_adj_* = 0.033). Males used treefall refuges more frequently in the wet season (dry/wet ratio = 0.18, *p_adj_* = 0.038) but increased their use of hidden microhabitats during the dry season, including under leaf litter (dry/wet ratio =1.37, *p_adj_* = 0.036) and under logs (dry/wet ratio =1.79, *p_adj_*= 0.041). *A. trivittata*, also showed seasonal shifts in microhabitat use independent of sex (GLMM: season*microhabitat: *χ2*(8)=31.92, *p* < 0.001; Fig. 3A, Table S9), with a greater use of hidden microhabitats during the dry season, including under leaf litter (dry/wet ratio =2.01, *p_adj_* < 0.001), inside tree fall refuges (dry/wet ratio =7.40, *p_adj_* < 0.001), and inside root cavities (dry/wet ratio =1.59, *p_adj_* = 0.034). In contrast, during the wet season, they showed greater use of exposed microhabitats, such as on top of the leaf litter (dry/wet ratio = 0.33, *p_adj_* < 0.001), and within leaf litter cavities (dry/wet ratio = 0.53, *p_adj_* = 0.007), classified here as potential clutch sites.

**Fig. 3.**
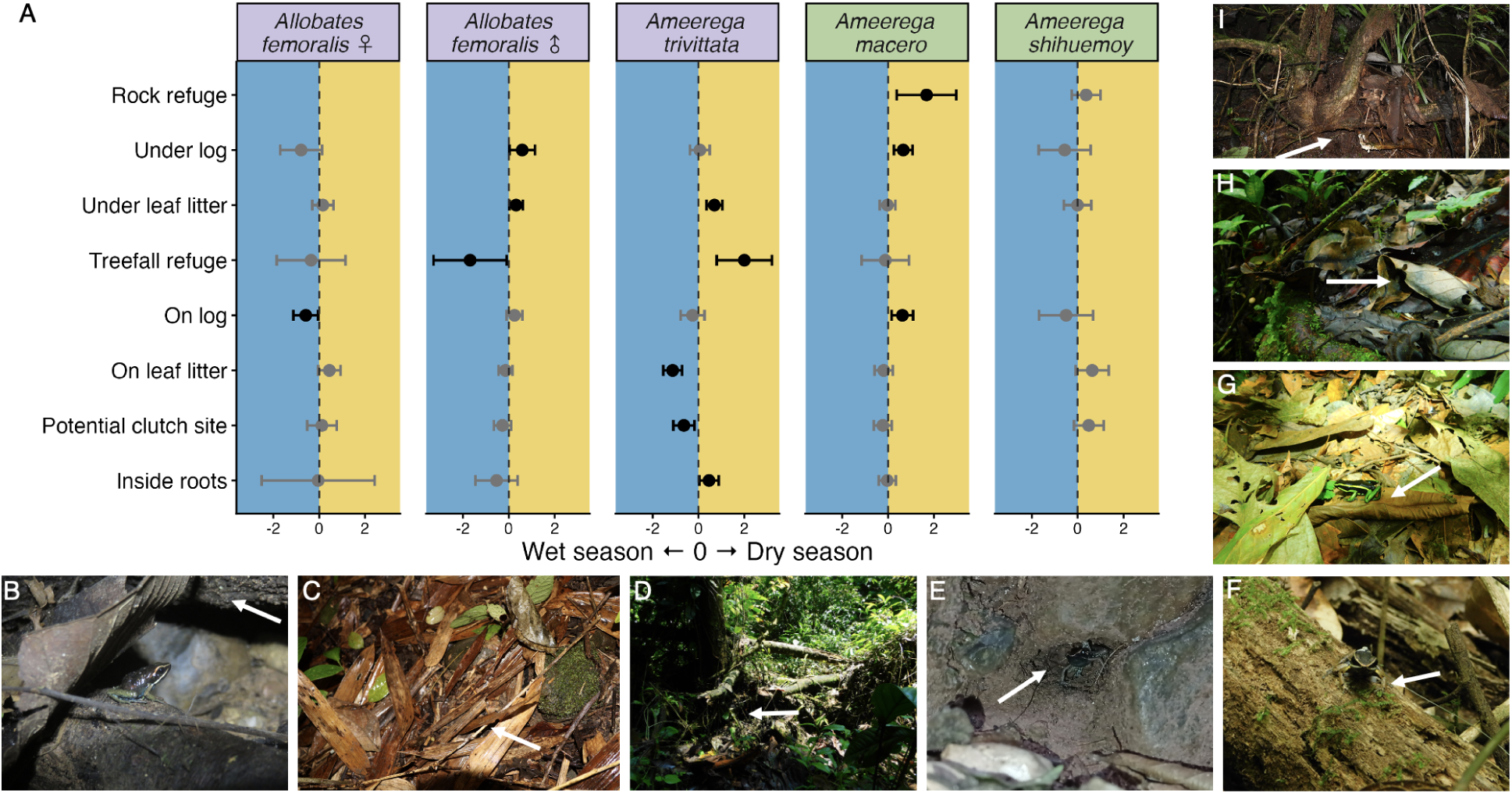
Seasonal shifts in microhabitat use across four poison frog species. (A) Pairwise contrasts (dry vs. wet) from generalized linear mixed models testing seasonal differences in microhabitat use for each species and sex. Points represent estimated log-odds ratios ± 95% CI. Values to the right of zero indicate higher use in the dry season; values to the left indicate higher use in the wet season. Black points denote significant contrasts (p < 0.05) and grey points non-significant contrasts. (B-I) Field photographs showing frogs using microhabitats (white arrows): (B) under log, (C) under leaf litter, (D) inside tree fall refuge, (E) inside rock refuge, (F) on log, (G) on leaf litter, (H) inside leaf litter cavity “potential clutch site”, (I) inside roots. Purple panels represent seasonal breeders and green panels represent year-round breeders. Sample sizes for panel A were: *A. femoralis* dry season (13 females, 23 males) and wet season (8 females, 26 males); *A. trivittata* dry (7 females, 11 males) and wet (8 females, 12 males); *A. macero* dry (11 females, 24 males) and wet (12 females, 23 males); and *A. shihuemoy* dry (11 females, 4 males) and wet (9 females, 10 males).

Among year-round breeders, shifts were species-specific. *Ameerega macero* showed significant seasonal shifts in microhabitat use (GLMM: season*microhabitat: *χ2*(9)=19.10, *p* = 0.024; Fig. 3A, Table S9), with greater use of log and rock-associated microhabitat during the dry season, including on top of logs (dry/wet ratio = 1.74, *p_adj_* = 0.046), under logs (dry/wet ratio =2.27, *p_adj_*< 0.001), and inside rock refuges (dry/wet ratio = 4.23, *p_adj_*= 0.035). In contrast, *A. shihuemoy* showed no seasonal variation in microhabitat use (GLMM: season*microhabitat: *χ2*(7) =6.35, *p* = 0.499).

### Water-borne testosterone and corticosterone release rates vary by species

Testosterone patterns varied among species. In the seasonal breeder *A. femoralis*, seasonal changes depended on sex (LM: season*sex: F_1,32_= 10.95, *p* = 0.002), while frog size had no effect (F_1,32_= 0.30, p = 0.587). Post hoc contrasts showed that *A. femoralis* females had significantly higher testosterone release rates in the dry season compared to the wet season (dry/wet ratio = 2.74, *p_adj_* < 0.001; Fig. 4A), whereas no seasonal difference was detected in males. No seasonal effects were detected in any of the other species (Supplementary Table S11). Additionally, testosterone release rates were not associated with space use in any species, with linear models revealing no significant effects of testosterone, sex, or their interaction on space use area (Supplementary Table S12).

**Fig. 4.**
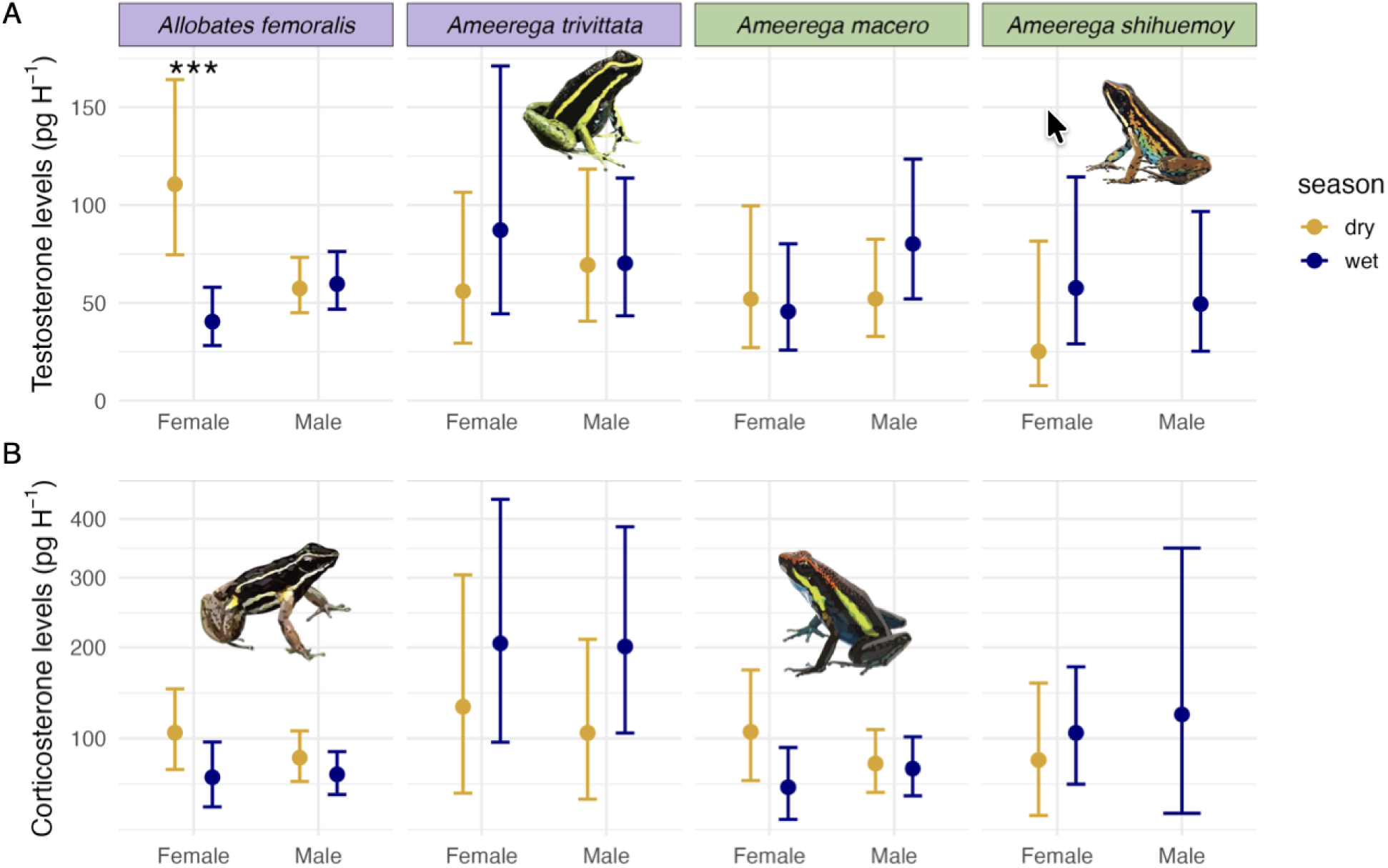
Seasonal variation in hormone release rates across four poison frog species. Estimated marginal means (±95% confidence intervals) of testosterone (A) and corticosterone (B) release rates (pg/h) for females and males of seasonal breeders (*A. femoralis* and *A. trivittata,* purple panel) and year-round breeders (*A. macero* and *A. shihuemoy,* green panel) across dry (yellow) and wet (blue) seasons. Points and error bars show back-transformed estimates from the log scale. Asterisks indicate significant seasonal differences within sex and species groups based on pairwise comparisons (*p* < 0.001***). Sample sizes for testosterone were: *A. femoralis* dry season (5 females, 13 males) and wet season (6 females, 13 males); *A. trivittata* dry (5 females, 7 males) and wet (6 females, 9 males); *A. macero* dry (5 females, 10 males) and wet (9 females, 12 males); and *A. shihuemoy* dry (3 females, 0 males) and wet (5 females, 8 males). Sample sizes for corticosterone were: *A. femoralis* dry season (6 females, 12 males) and wet season (6 females, 14 males); *A. trivittata* dry (4 females, 8 males) and wet (9 females, 8 males); *A. macero* dry (5 females, 12 males) and wet (10 females, 14 males); and *A. shihuemoy* dry (6 females, 0 males) and wet (7 females, 3 males).

Corticosterone patterns also varied among species but showed no significant seasonal variation in either sex across all four species (Fig. 4B; Supplementary Table S13).

### Limited chemical defense variability across seasons

Across species, we detected 26 alkaloids assigned to eight structural families, with one unclassified and one unknown alkaloid (Supplementary Table S2). No alkaloids were detected in the seasonal breeder *A. femoralis*. Detection frequency varied strongly among species and seasons (Fig. S4). *Ameerega trivittata* exhibited the highest alkaloid richness, with 20 alkaloids detected during the wet season and 19 during the dry season. *A. macero* and *A. shihuemoy* exhibited lower richness overall, with 8 and 7 alkaloids detected during the wet season and 13 and 11 during the dry season, respectively. Mean alkaloid abundance varied among individuals within species and seasons (Fig.5). Species-specific PERMANOVA analyses revealed no evidence of seasonal differences in overall alkaloid composition within species (*A. trivittata:* p = 0.212; *A. macero*: p = 0.361; *A. shihuemoy*: p = 0.413; Supplementary Table S14). Similarly, Wilcoxon rank-sum tests revealed limited evidence of seasonal differences in alkaloid families within species (Supplementary Table S15).

**Fig. 5.**
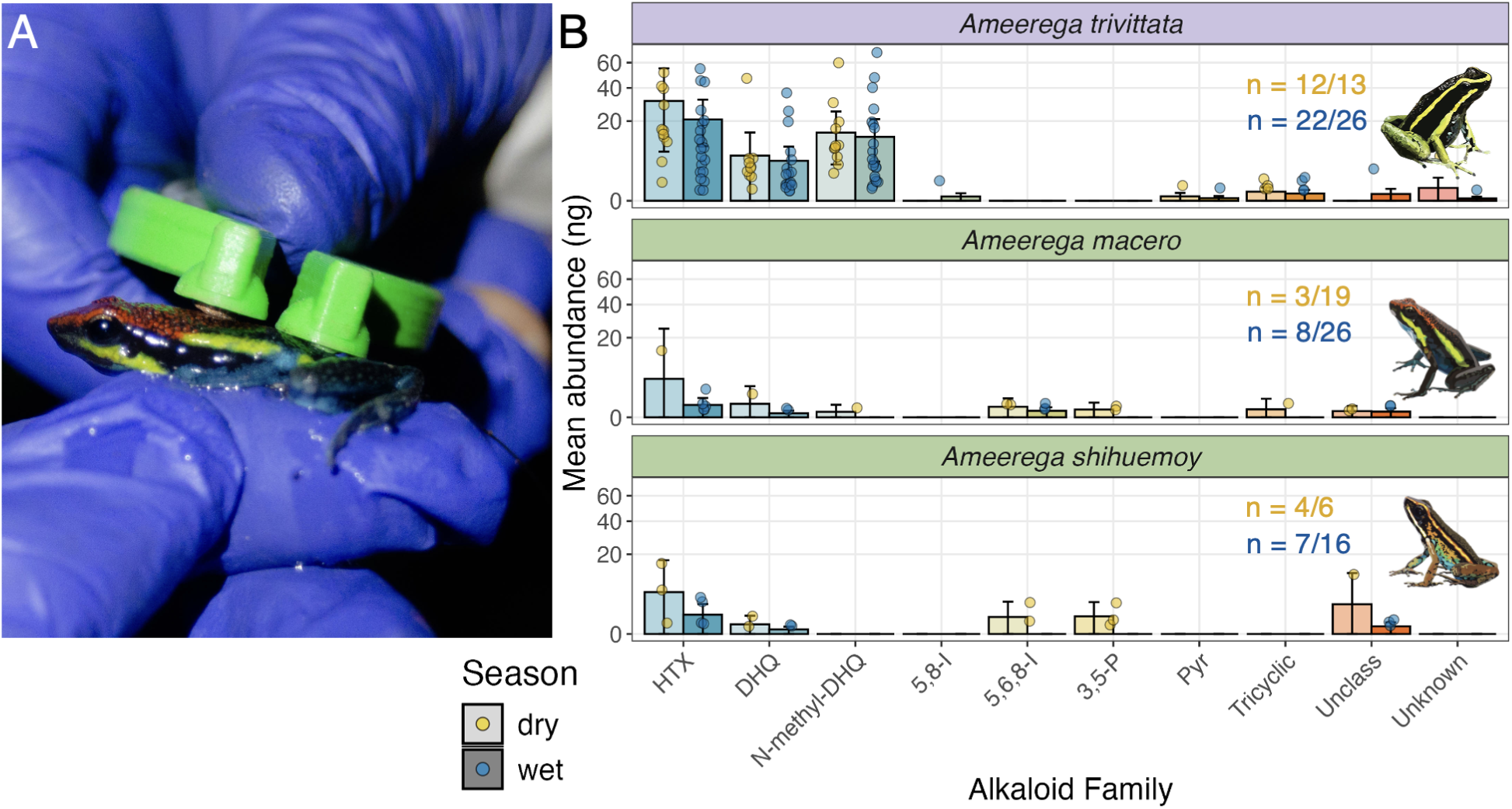
Seasonal variation in poison frog chemical defenses. (A) Field photograph illustrating electrical stimulation used to induce skin alkaloid release with a Ranavolt device. (B) Mean abundance (ng) of alkaloid families within each species–season combination (sample sizes indicated as samples with detectable alkaloids/total sampled). Error bars indicate 95% confidence intervals, and points represent individual frogs with detectable alkaloids. Alkaloid abundance is shown on a square-root scale. Alkaloid families include histrionicotoxins (HTX), decahydroquinolines (DHQ), N-methyl decahydroquinolines (N-methyl-DHQ), 5,8-disubstituted indolizidines (5,8-I), 5,6,8-trisubstituted indolizidines (5,6,8-I), 3,5-disubstituted pyrrolizidines (3,5-P), piperidines (Pyr), tricyclic alkaloids, unclassified alkaloids, and unknown alkaloids. No alkaloids were detected in the seasonal breeder *Allobates femoralis*.

## Discussion

In this study, we integrated fine-scale spatial behavior, microhabitat use, hormone release rates, and chemical defenses across four poison frog species with contrasting breeding strategies. We found species-specific behavioral responses to seasonal variation across both space use area and microhabitat use. Seasonal breeders exhibited pronounced seasonal changes in space use, whereas year-round breeders maintained relatively stable space use across seasons. At the microhabitat level, *A. femoralis*, *A. trivittata*, and *A. macero* shifted substrate preferences seasonally, whereas *A. shihuemoy* maintained broad use of multiple substrates across seasons. These behavioral strategies occurred without strong seasonal shifts in testosterone, corticosterone, or chemical defenses. Thus, these results suggest that poison frogs may cope with seasonal variability primarily through behavioral buffering mechanisms, supporting a potential decoupling between behavioral and some physiological responses to environmental change.

### Reproductive strategy shapes sex-specific seasonal variation in space use area

Seasonal effects on space use were species- and sex-specific, likely reflecting seasonal changes in reproductive activity and environmentally driven variation in movement. In both seasonal breeders, *A. femoralis* and *A. trivittata*, males expanded their space use during the wet season, which coincides with their peak reproductive activity (Summers and Tumulty 2014). This expansion excluded movement associated with transporting tadpoles, because poison frogs typically carry their offspring to water bodies outside their daily movement range (Pašukonis et al. 2019). During the dry season, males reduce space use, likely reflecting reduced breeding behaviors and limited movement in response to conditions that increase the risk of overheating and dehydration. Many researchers have documented comparable seasonal contractions in space use during unfavorable periods across a range of ectotherms, including tortoises (Nowakowski et al. 2020), lizards (Ariano-Sánchez et al. 2020), caimans (Mascarenhas-Junior et al. 2024), and amphibians (Watson et al. 2003), suggesting a common strategy to seasonal variation across terrestrial vertebrate ectotherms. Recaptured individuals showed variable degrees of overlap in locations used between dry and wet seasons, indicating that shifts in space use were not solely driven by expansion or contraction of used areas, but also by relocation of those areas. In *A. femoralis*, both males and females exhibited low area overlap with only minor shifts in activity centers. In contrast, *A. trivittata* showed complete relocation of space use in both sexes, accompanied by large centroid shifts, suggesting low inter-seasonal side fidelity.

While males of both seasonal breeders expanded their space use area during the wet season, females of *A. femoralis* showed the opposite pattern, exhibiting larger space use during the dry season. Rather than tracking reproductive opportunities, females may be more strongly driven by foraging requirements during the dry season, when arthropod biomass declines (Newell et al. 2023; Pedroso-Santos et al. 2024). Similar foraging-driven expansions in space use have been documented in other taxa (Natusch et al. 2021). In contrast, males did not show a comparable dry-season expansion. One possible explanation is that territorial behavior constrains short-term movement even under changing environmental conditions. We recorded similar numbers of aggressive encounters across seasons (wet season: n = 4; dry season: n = 3), suggesting continued territorial activity during the dry season. However, previous work has shown that male *A. femoralis* abandon territories between reproductive seasons and subsequently re-negotiate them when reproduction resumes (Ringler et al. 2009), indicating that movement dynamics may be different depending on the temporal scale considered. These findings suggest that males and females may experience different selective pressures imposed by the same seasonal environmental challenges. Accordingly, comparative work across ectotherms shows greater thermal tolerance plasticity in females than males (Pottier et al. 2021), underscoring the importance of analyzing sexes separately.

Year-round breeders *A. macero* and *A. shihuemoy* showed no seasonal differences in space use area, suggesting alternative strategies for accessing reproductive resources year-round. Unlike the seasonal breeders, these species rely more on stream margins during the dry season, which may provide relatively stable breeding resources across seasons. Similar seasonal stability in space use has been described in other vertebrates (e.g., mammals: Mandl et al. (2018) and birds: Odom et al. (2019)). This relative stability in space use is consistent with the limited centroid shifts observed in recaptured individuals, suggesting that individuals maintain broadly similar activity centers across seasons.

An important limitation of our study concerns the classification of *A. shihuemoy* as a year-round breeder. Although this classification was based on previous natural history observations, our own observations of reproduction did not clearly support continuous breeding across seasons. In particular, we detected more reproductive events during the dry season than the wet season, introducing uncertainty regarding the species’ reproductive timing. Although seasonal differences in detectability may partially contribute to this pattern, additional long-term natural history and reproductive monitoring will be necessary to determine whether this species truly reproduces continuously throughout the year.

Complementary analyses showed species-specific relationships between daily movement and environmental conditions, further suggesting that climatic factors influence movement differently among species. In the seasonal breeder *A. femoralis*, increased rainfall was associated with greater daily movement, consistent with heightened activity during the wet season when reproductive opportunities peak and dehydration constraints are reduced. Increased movement during the wetter period may facilitate reproductive behaviors and access to tadpole deposition sites, while simultaneously lowering the physiological costs associated with movement in humid conditions. This pattern suggests that rainfall may directly influence the balance between reproductive activity and environmental constraints in this species. In contrast, movement in *A. trivittata* and *A. macero* was more strongly associated with temperature, suggesting that thermal conditions may play a larger role in modulating activity in these species. These differences indicate that climatic drivers of movement are not uniform across species, even within similar habitats. Notably, *A. shihuemoy* showed no detectable relationship between movement and environmental variables, which may reflect limited statistical power. Together, these findings suggest that seasonal changes in space use are not only determined by shifts in area or location, but also by species-specific movement responses to environmental conditions.

### Microhabitat use as a behavioral buffering mechanism against seasonal variability

A growing body of research highlights the key role of behavioral buffering through microhabitat selection in mediating the effects of climate and land use change on frogs (González-del-Pliego et al. 2020; Klinges et al. 2024; Riddell et al. 2026). We found seasonal shifts in microhabitat use that suggest that fine-scale microhabitat selection acts as a behavioral buffering mechanism against environmental stress. This buffering operates through behavioral thermoregulation and water loss regulation, allowing ectotherms to reduce exposure to unfavorable environmental conditions (Kearney et al. 2009; Rozen-Rechels et al. 2019). Such mechanisms are particularly important for amphibians because their permeable skin makes them highly susceptible to overheating and dehydrating.

In line with a buffering role, seasonal breeders shifted toward hidden, humid microhabitats during the dry season, including leaf litter, fallen logs, treefall refuges, and root cavities. Similar seasonal shifts have been documented in both temperate and tropical frogs (Eterovick et al. 2010; Brandão et al. 2020; Park et al. 2024). These microhabitats buffer temperature extremes and reduce water loss, minimizing exposure to harsh climatic conditions (Scheffers et al. 2014). During the dry season, the reduced use of exposed calling sites likely reflects individuals investing more energy in survival than reproduction under environmental constraints (Reniers et al. 2015). Responses among year-round breeders were more species-specific. Although no strong microhabitat preference was observed during the wet season, *A. macero* shifted toward exposed calling sites during the dry season, whereas *A. shihuemoy* maintained relatively consistent use of multiple microhabitats across seasons. The proximity of their home ranges to streams likely provides a continuous humid buffer because habitats closer to stream margins typically maintain higher humidity and reduced thermal variability than surrounding forest (Hoffmann et al. 2021; Aguilar et al. 2025; Cecala et al. 2025). The broad microhabitat use observed in *A. shihuemoy* reflects a more generalized microhabitat strategy. Together, these findings suggest that poison frogs buffer seasonal variability through different behavioral mechanisms, including seasonal shifts in substrate preference or stable use of diverse microhabitats across seasons.

Together, these findings emphasize the importance of microhabitat structure to buffer environmental stress and support our findings that poison frogs primarily rely on behavioral flexibility to buffer seasonal variability. With the expected unpredictability of tropical rainfall, effective microhabitat buffering will become increasingly important. However, while behavioral flexibility can mitigate short-term climatic variability (Woods et al. 2015; Sears et al. 2016), its effectiveness depends on the long-term availability of these microhabitats (González-del-Pliego et al. 2020).

### Behavioral responses may be partially decoupled from hormonal seasonality

Contrary to our predictions, the pronounced seasonal behavioral changes observed in seasonal breeders were inconsistent with seasonal shifts in corticosterone or testosterone release rates. This pattern suggests that substantial behavioral adjustments to seasonal environmental variation may occur without corresponding changes in water-borne hormone levels. Such partial decoupling between behavioral and hormonal seasonality contrasts with findings in other taxa (Wingfield et al. 1990; Landys et al. 2010), including amphibians and reptiles (Eikenaar et al. 2012; Lind et al. 2018). In vertebrates, corticosterone and testosterone regulate life-history trade-offs associated with metabolic regulation, responses to environmental variation, and reproductive effort (Wingfield et al. 1998; Moore and Jessop 2003; Denver 2009b; Husak et al. 2021). Our results show limited evidence for pronounced seasonal hormonal shifts, with the exception of *A. femoralis* females. Instead, both seasonal and year-round breeders seem to respond to environmental variation primarily through behavioral adjustments rather than strong hormonal modulation.

Corticosterone release rates remained stable across seasons. In tropical environments, dry periods are typically characterized by reduced resource availability, increased dehydration risk, and constraints on reproduction (Gesquiere et al. 2008). Such constraints are often expected to elevate glucocorticoid levels, which are adaptive responses to increased energy and water demands (Maute et al. 2013; Moeller et al. 2023). However, in some species, glucocorticoids are higher during the breeding season to meet the energy demands of testosterone-regulated reproductive behavior. (Eikenaar et al. 2012; Crocker-Buta and Leary 2018). Thus, elevated glucocorticoids likely reflect life energetic allocation rather than direct responses to environmental stressors. When organisms can behaviorally buffer challenging conditions through microhabitat selection or activity shifts, physiological shifts may remain limited (Kearney et al. 2009; Dezetter et al. 2023). In this context, the absence of a seasonal shift in corticosterone is consistent with a scenario of partial behavioral buffering, although this interpretation should be made cautiously given the inherent variability of water-borne hormone measures.

Testosterone release rates remained stable across seasons, except for *A. femoralis* females, which showed higher levels in the dry season. This increase may be associated with changes in social dynamics under less favorable environmental conditions. Elevated female testosterone has been associated with increased aggression and reduced reproductive behaviors across vertebrates (Veiga and Polo 2008; French et al. 2013; Enbody et al. 2018; Crespi et al. 2025). If similar mechanisms are acting in *A. femoralis* females, higher dry-season testosterone could reduce mating receptivity when environmental conditions are constrained; however, alternative explanations remain possible, including a role in supporting increased demands during the dry season. For example, if elevated testosterone reflects increased aggression or reduced mating receptivity, dry season females may show changes in social behavior, which needs further experimental testing.

If testosterone contributes to seasonal spatial behavior, individuals with higher testosterone release rates would also be expected to exhibit larger space use areas. However, despite clear seasonal differences in space use, we found no evidence of an association between testosterone release rates and space use in any species. This result contrasts with studies in frogs (Pašukonis et al. 2022) and mammals (Burkitt et al. 2007; Schulz and Korz 2010), where testosterone has been linked to spatial activity. Together, these findings strengthen the interpretation that seasonal behavioral responses in these poison frogs may be at least partially decoupled from circulating testosterone levels measured via water-borne sampling. Instead, behavioral adjustments to seasonal variability may rely more strongly on immediate ecological responses or on neuroendocrine mechanisms not captured by baseline water-borne hormone measurements.

Some methodological limitations may also contribute to the absence of strong seasonal hormonal differences. Water-borne hormone measurements integrate hormone release over the standardized one-hour sampling period and therefore represent an integrated estimate of hormone secretion influenced by recent physiological and environmental conditions. As a result, variation among individuals may reflect short-term variation in hormone release dynamics associated with behavioral context, time of day, or recent environmental exposure. Although we recorded behavior immediately prior to capture, observations were brief and most individuals were classified as perching, limiting our ability to assess whether immediate behavioral state contributed to variation in hormone release rates. Future studies incorporating longer pre-capture observation periods would be valuable for disentangling short-term behavioral effects from general seasonal patterns. Such variability can obscure seasonal patterns when measurements are averaged across individuals. Moreover, frog hormone levels can change rapidly in response to social or environmental stimuli, sometimes within short time scales (Rodríguez et al. 2022).

Further limitations relate to detectability and biological variability in water-borne hormone assays. Water-borne measurements inherently vary among individuals due to differences in secretion rates, requiring larger sample sizes per group to reliably detect effects (Gabor et al. 2016, 2018). Although our sample sizes are comparable to other amphibian studies (Rodríguez et al. 2022), this variability may reduce power to detect seasonal endocrine changes. Therefore, non-significant hormonal patterns should be interpreted cautiously, as they may reflect methodological and biological variability rather than true absence of seasonal hormonal variation. Additionally, behavioral responses may depend on other endocrine mechanisms not measured here, such as variation in hormone receptor density or receptor binding in target tissues (Baugh et al. 2018), which could be explored in future studies. Despite these limitations, the repeated observation of seasonal behavioral changes alongside limited endocrine variation suggests that behavioral buffering may play a larger role in responses to seasonal variability in tropical amphibians.

### Limited seasonal variation in chemical defenses

We found limited evidence for seasonal changes in poison frog chemical defenses. These results are partially in line with the findings of Basham et al. (2021), who reported no seasonal difference in alkaloid richness in the poison frog *Andinobates fulguritus*. Similarly, Moskowitz et al. (2018) documented seasonal diet shifts in the poison frog *Mantella laevigata,* yet their overall assortment of identified alkaloids was similar across seasons, suggesting that alkaloid profiles may be relatively robust to short-term environmental variation. In contrast, Saporito et al. (2006) found significant seasonal alkaloid variation in *Oopagha pumilio*, although spatial variation in prey availability may have contributed to these differences. These comparisons suggest that seasonal environmental variation can influence poison frog chemical defenses in multiple ways, depending on the ecological context.

Our results align with previous studies showing strong species differences in frog chemical defenses. Most alkaloids detected in *A. trivittata* and *A. macero* have been previously reported. For *A. shihuemoy*, this study provides the first assessment of alkaloid composition. Consistent with previous research on *Ameerega*, histrionicotoxins were the most abundant alkaloid family across species (Daly et al. 2009; Albuquerque-Pinna et al. 2024). In addition, the occurrence of both decahydroquinolines (DHQs) and N-methyl decahydroquinolines (N-methyl DHQs) in *A. macero* and *A. trivittata* is consistent with previous evidence of metabolic transformation of DHQs into N-methyl DHQs in several *Ameerega* species (Daly et al. 2009) This metabolic conversion has also been reported in other dendrobatid genera, including *Adelphobates* (Jeckel et al. 2022) and *Ranitomeya* (Stuckert et al. 2014). In contrast, N-methyl DHQs was not detected in *A. shihuemoy*, although it remains unclear whether they are truly absent or occurred below the detection limit of our extraction and analytical methods.

The absence of detectable alkaloids in *A. femoralis* should be interpreted cautiously. Although this result limited the inclusion of this species in seasonal comparisons of chemical defenses, previous studies using full-skin GC-MS extractions have similarly reported either undetectable or extremely low alkaloid quantities in wild *A. femoralis* populations (e.g., (Daly et al. 1987; Saporito and Grant 2018; Tarvin et al. 2024; Jeckel et al. 2026)). Our swab-based collection method is less complete than full-skin extraction methods, raising the possibility that alkaloids were present at concentrations below the detection threshold of our analyses. However, methodological limitations alone are unlikely to fully explain these results, given that low alkaloid abundance has also been documented using more exhaustive sampling approaches. Experimental studies further demonstrate that *A. femoralis* is capable of sequestering alkaloids when exposed to alkaloid-containing diets (Sanchez et al. 2019; Caty et al. 2025), suggesting that alkaloid expression in *A. femoralis* may be naturally low or highly variable across populations.

Importantly, our ability to detect seasonal effects was constrained in some species by the low number of individuals with detectable alkaloids during the dry season, particularly in *A. macero* and *A. shihuemoy*. This substantially reduced statistical power and increased uncertainty in estimates of seasonal alkaloid abundance and composition, as reflected by the broad confidence intervals observed in some species–season combinations. We therefore interpret the absence of significant seasonal differences cautiously and cannot rule out the possibility that subtle seasonal changes in chemical defenses were not detected. Nevertheless, the available data suggest that interspecific differences in alkaloid composition may be stronger than seasonal differences within species.

## Conclusions

Our results reveal two behavioral strategies through which poison frogs cope with rainfall seasonality. Seasonal breeders appear to rely on predictable rainfall patterns by concentrating reproductive investment in the wet season and reducing space use while selecting microrefugia during the dry season, minimizing exposure to unfavorable conditions. Year-round breeders maintain reproductive activity across seasons, by relying on microhabitats with persisting humid refugia, particularly closer to streams, and maintaining the same spatial use. These strategies have distinct implications for resilience under climate change. For seasonal breeders, the primary risk is disruption of the seasonal predictability, as climate projections for tropical regions consistently forecast alterations in rainfall timing and intensity. For year-round breeders, the increased uncertainty of rainfall patterns may surpass the buffering capacity of species that depend on microhabitat refugia, threatening adaptations that enable year-round reproduction. By revealing how frogs currently cope with seasonal variability, this study provides insights into how species with contrasting life-history traits may respond as climate change continues altering the conditions under which these strategies evolved.

## Author contributions

SJSR: conceptualization, methodology, field data collection, data curation, sample processing, data analysis, writing—original draft, writing—review & editing. LAO: conceptualization, funding acquisition, methodology, writing—review & editing. AP: conceptualization, methodology, data collection, writing—review & editing. MG: methodology, sample processing, writing—review & editing, CR: methodology, writing—review & editing. All other co-authors have participated in field data collection, data curation, and writing—review & editing.

## Animal ethics

This research was conducted in accordance with appropriate research permits (RD N° D000133-2022-MIDAGRI-SERFOR-DGGSPFFS-DGSPFS, authorization code N° AUT-IFS-2022-084 and N° D000049-2023-MIDAGRI-SERFOR-DGGSPFFS-DGSPFS) and CITES permits (N° 005157/SP and N° 005871/SP) issued by the local authorities, and was approved by the Institutional Animal Care and Use Committee of Stanford University (protocol ID 33211 and 33839 issued to LAO).

## Acknowledgements

We thank Dr. Rodolfo Dirzo for his valuable guidance on the methodology and analytical approach, and for his insightful comments during the revision of the manuscript. We thank Prof. Robyn B. Lockwood for providing suggestions that improved the clarity and readability of the manuscript. We thank the NGOS Amazon Conservation and Crees Foundation for their hospitality during fieldwork. We thank all the volunteers in both organizations who help us with information on frog observations. We acknowledge with sadness the passing of Jhony Oswaldo Turizo and express our sincere gratitude for his design and construction of the Ranavolt. This device was fundamental to implementing a non-invasive method for collecting alkaloid secretions from the poison frogs studied and was critical to the successful completion of this work. His contribution remains an integral part of this research, and we are deeply appreciative of his collaboration and dedicated assistance throughout the development of this project. We are also grateful to the editor and anonymous reviewers for their constructive feedback, which substantially improved the clarity and quality of this manuscript.

