## Supplemental Files for "Behavioral responses buffer seasonal variation more strongly than endocrine and chemical responses in poison frogs"

**Supplementary Methods**

**Supplementary Methods S1: Environmental variables**

Temperature was characterized as mean daily temperature, and precipitation was characterized using three complementary metrics: rainfall probability (frequency of rainy days), rainfall intensity (mean rainfall on wet days), and daily precipitation (total rainfall per day). Daily precipitation for the Manu Learning Centre is shown in Fig. 1C, with comparable environmental data for Los Amigos Biological Station shown in Supplementary Fig. S1.

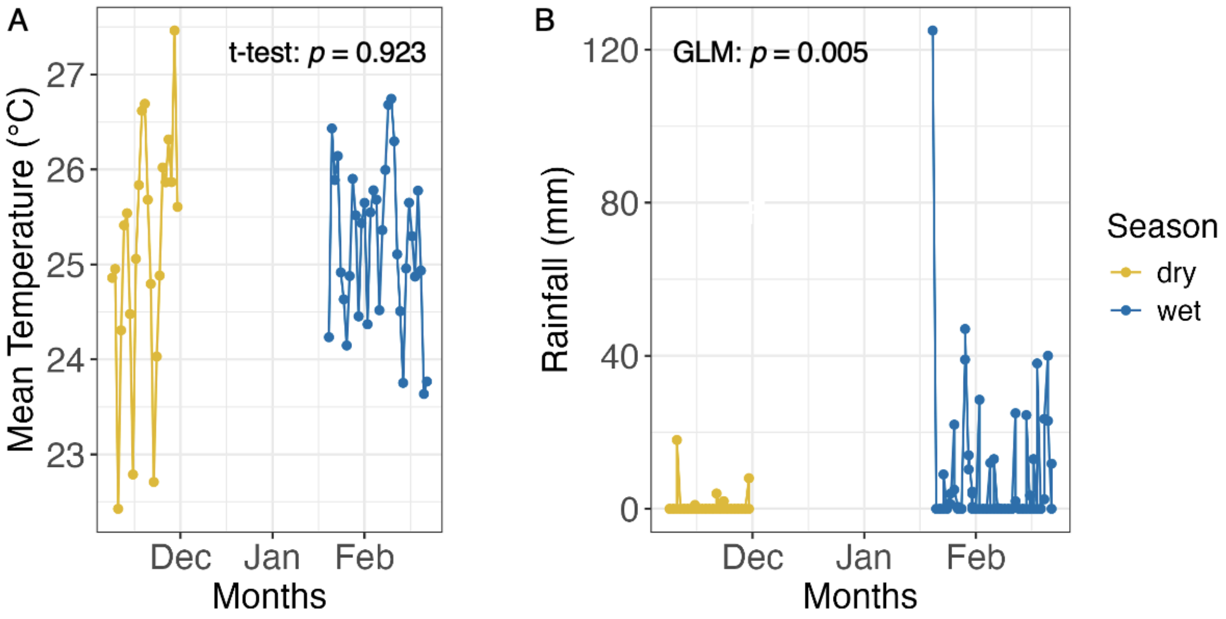


**Fig. S1** (A) Mean temperature and (B) daily precipitation during the dry and wet seasons at Los Amigos Research Station, where *A. trivittata* was studied.

**Supplementary Methods S2: Study species**

In this study, we focus on four poison frogs with contrasting breeding duration strategies: two seasonal breeders (*A. femoralis* and *A. trivittata*) and two year-round breeders (*A. macero* and *A. shihuemoy*).

***Allobates femoralis*** (brilliant-thighed poison frog) is widely distributed throughout South America and found in tropical lowland forests up to 1000 m asl. It is a diurnal species with peak activities between 06:00 and 10:00 hours and between 14:00 and 18:00 hours (Ringler et al. 2009). Males exhibit strong territoriality during the breeding season, beginning at the onset of the rainy season (Kaefer et al. 2012), while territories are abandoned during the dry season (Ringler et al. 2009), marking their reproductive activity concentrated during the wet season.

***Ameerega trivittata*** (three-striped poison frog) is widely distributed throughout South America and found in tropical rainforests up to 680 m asl. It is a crepuscular and diurnal species with peak activities between 04:40 and 09:00 hours (Roithmair 1994) and between 14:00 and 18:00 hours (personal observation). They are one of the largest dendrobatid frogs with the longest homing behavior recorded (~ 800 m; Pašukonis et al. 2018). They have been observed transporting up to 41 tadpoles to pools and creeks, usually outside their territory (Neu et al. 2016; Pašukonis et al. 2018). Although frogs can be seen throughout the year, calling and reproducing activity ceases during the dry season (Roithmair 1994).

***Ameerega macero*** (Manu poison frog) is widely distributed throughout Southeastern Peru, extending to the western part of Brazil, at elevations ranging from 300 to 550 m asl (Rodríguez and Myers 1993). It displays a crepuscular pattern, with higher activity observed between 05:00 and 07:00 hours and again between 16:00 and 18:00 hours. While the ecology of this species remains poorly understood, we observed adults calling and transporting tadpoles throughout the year during four years of prior fieldwork in the study area. These long-term observations provide evidence for a year-round breeding strategy.

***Ameerega shihuemoy*** (Amarakaeri poison frog) is an endemic species from the Manu Biosphere Reserve, found at elevations between 340 and 600 m asl. This species displays a crepuscular pattern, with higher activity observed between 05:30 and 09:00 hours, and between 16:00 and 18:00 hours. This species is characterized by its year-round breeding pattern. In the dry season, it selects forest streams as its breeding grounds, utilizing the small water bodies formed along the stream margins, facilitated by the slow water flow. When the rainy season arrives, this species moves towards the forest interior to deposit tadpoles in temporary pools within the forest (Serrano-Rojas et al. 2017).

During the study period, seasonal breeders displayed uneven seasonal distribution of reproductive observations (egg deposition and tadpole transport), with most events occurring in the wet season (*A. femoralis*: dry = 5, wet = 9; *A. trivittata*: dry = 4, wet = 11). Year-round breeders reproduced in both seasons, with observations evenly distributed in *A. macero* (dry = 6, wet = 6), whereas *A. shihuemoy* showed more reproductive activity in the dry season (dry = 8, wet = 2). Although our sample sizes were too small for formal statistical tests, the repeated reproductive behaviors across seasons in year-round breeders, the skewed seasonal distribution in seasonal breeders, and prior natural history observations support this classification. The apparent skew toward more reproductive behaviors in *A. shihuemoy* during the dry season may reflect sampling bias: tadpole-carrying adults are easier to observe along open streams in the dry season, whereas in the wet season, they are more difficult to detect in the forest interior (Supplementary Fig. S2).

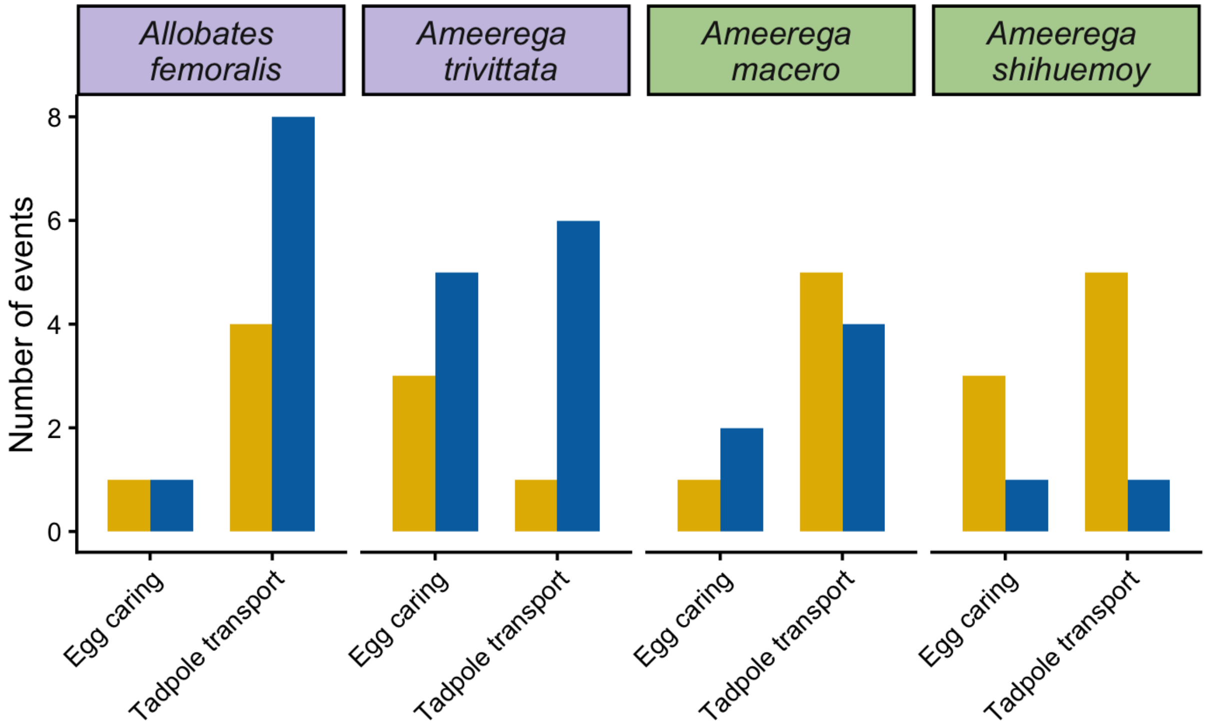


**Fig. S2** Number of parental care events recorded during the dry and wet seasons for four poison frog species. Bars represent the total number of observed egg caring and tadpole transport events per species and season (dry = yellow; wet = blue). Purple panels represent season breeders, and green panels represent year-round breeders.

**Supplementary Methods S3: Telemetry method details**

For harmonic direction finding (HDF), we used a handheld transceiver R9 (RECCO detector 99B, Lindigö, Sweden) to detect miniature transponders attached to frogs with a silicone waistband (Gourret et al. 2011; Beck et al. 2017). The tags weighed ~0.09 g and constituted < 10% of the body weight of an adult *A. femoralis* (mass range: 1.0–2.25 g), *A. macero* (1.12–2.66 g), and < 11% for the smallest *A. shihuemoy* (0.79–1.41 g). For radio-tracking with very high frequency (VHF) transmitters, *A. trivittata* was equipped with miniature very high frequency transmitters (NTF-2-1 nanotag freshwater, Lotek Wireless Inc.). Frogs were located using a portable radio-tracking receiver (Biotracker30 receiver) and a flexible Yagi antenna (Liteflex Yagi). The tags weighed 0.30–0.35 g and constituted <10% of adult *A. trivittata* (3.45–7.36 g).

To quantify space use, we tagged 36 and 34 *A. femoralis* individuals in the dry and wet seasons, respectively; 18 and 20 A. trivittata; 35 *A. macero* in both seasons; and 15 and 19 *A. shihuemoy* in the dry and wet seasons, respectively. In total, we localized individuals 13,594 times and tracked them for periods ranging from 2 to 22 days per individual.

**Supplementary Methods S4: Water-borne hormone collection and quantification**

We collected water-borne hormone samples following a non-invasive amphibian hormone sampling method originally developed by Gabor et al. (2013) and subsequently adapted for other frog species, including poison frogs (Baugh et al. 2018; Rodríguez et al. 2022). We captured all individuals in the afternoon (between 15:00 and 18:00 h) to control for potential diel variation in hormonal changes throughout the day. We minimized disturbance during capture time to reduce as much as possible handling that could alter hormonal levels. Frogs were carefully grasped by their tracking antenna and immediately placed into clean, transparent ziplock bags, avoiding the inclusion of soil or leaf litter that could contaminate the sample.

Each frog was then transferred to a glass container (14 × 9 × 5 cm) containing 40 mL of distilled water, within approximately 1–3 min of initial capture. The container was covered with an upside-down bucket to create a dark environment and minimize further disturbance. Frogs remained in the container for one hour to allow hormones released through the skin and urine to accumulate in the water. After this period, we removed the bucket and glass container lid and allowed each frog to jump freely back into the forest while still retaining the tracking tag. The 40 mL of water was then collected as the hormone sample for subsequent analysis. We immediately filtered each sample using a 20 mL sterile syringe connected to an individual C18 cartridge (SPE, Sep-Pak C18 Plus, 360 mg sorbent, 55–105 µm particle size, #WAT020515, Waters Corp., Milford, MA, USA) at a flow rate of approximately 10 mL/min. Cartridges were used directly, following previous amphibian water-borne hormone studies (Pašukonis et al. 2022). Hormones retained in the cartridge were eluted with 4 mL of absolute ethanol into 5 mL glass vials. As hormone samples were collected across multiple field seasons, the time between sample collection and laboratory assays varied from 8 to 12 months. To reduce degradation of the hormones over time, we stored the samples at 4°C in the field and at -80°C in the laboratory until further processing. Ethanol preservation has been shown to effectively maintain steroid hormone concentrations in plasma during field storage prior to freezing, making this approach feasible for remote sampling (Goymann et al. 2007). To account for potential ethanol evaporation during storage, all samples were dried under nitrogen gas ($N_2$) at 37 °C before assay and reconstituted in 4 mL of 100% ethanol.

We quantified corticosterone and testosterone using commercial enzyme immunoassay kits (corticosterone: ADI-900-097; testosterone: ADI-900-065, Enzo Life Sciences, Farmingdale, NY, USA). We transferred 1ml of each reconstituted sample into a 1.5 ml Eppendorf tube and dried them in a SpeedVac concentrator at 37 °C. We then resuspended all samples with 250 µL of the assay buffer (provided in the kit) and incubated them overnight at 6 °C. Samples were brought to room temperature and shaken at 500 rpm for 1 h before the assay. All samples were plated in duplicate, and the assays were conducted according to the manufacturer's protocols. We read plates at 405 nm with correction between 570 and 590 nm, using a microplate reader (Synergy H1, BioTek Instruments, Winooski, VT, USA). We calculated hormone concentrations (pg/mL) using a four-parameter logistic curve in the software Gen5 (version 3.05, BioTek Instruments, Winooski, VT, USA).

To estimate water-borne release rates, we multiplied hormone concentrations obtained from the enzyme immunoassays by the final assay reconstitution volume (250 uL) and by the extraction dilution factor (4x). Values were expressed as hormone release rates (pg/h) because frogs were held in water for a standardized collection period of 1 h. We excluded all samples with a coefficient of variation (CV) > 20% from later analyses.

**Supplementary Methods S5: Alkaloid collection method details**

We collected samples using rayon regular tip swabs (Puritan Medical Products, Guilford, ME, USA, cat no. 25-806 1PR). Prior to sampling, all swabs were pre-treated by submerging them in beakers with approximately 15 mL of HPLC-grade methanol. We stirred each swab in methanol for one minute, discarded the residual, and repeated this washing procedure twice. We then inverted the swabs, placed them in an Erlenmeyer flask (to preclude contact with the swab tips), and left them to air-dry for at least 8 hours in a closed fume hood. Once dried, we stored swabs inside their original packaging and kept them in hermetic plastic bags until field sampling.

We captured all individuals one day after hormonal sampling, between 10:00 and 18:00 h, and followed the same capture protocol used during hormone sampling to minimize disturbance. We induced alkaloid secretion for three minutes using a Ranavolt set to 100 Hz and a duty cycle of 30% for all species. Voltage was set to 0.98 V for *A. shihuemoy* and 1.01 V for the other three species. Voltage was adjusted based on each frog’s behavioral response, ensuring visible muscle contraction in the body while avoiding the involuntary leg extension (Gonzalez 2021). Before stimulation, we rinsed the frog’s dorsum with a few drops of distilled water and placed the electrodes on the dorsum and back legs for three consecutive minutes. Immediately after stimulation, we swabbed the central, right, and left lateral areas of the dorsum and the dorsum of the front and hind legs, making 10 swabbing motions per area using the tip of a pre-cleaned swab. We cut each swab tip and placed it into a 2.0 mL empty glass vial (Agilent Technologies, Santa Clara, CA, USA; cat no. 5182-0714) and stored all dry samples at 4°C for up to four months, followed by storage at -80 upon return to Stanford until further processing. As alkaloid samples were collected across multiple field seasons, the time between sample collection and laboratory assays varied from 12 to 18 months. After sampling, we immediately rinsed the frogs with distilled water to restore body hydration and check the welfare of the animal, observing they were breathing normally and responding to mechanical stimuli. After confirming normal activity, we removed all tracking tags from all individuals at their site of last recapture immediately following alkaloid sampling. No adverse effects were observed.

**Supplementary Methods S6: Alkaloid extraction and annotation method details**

For alkaloid extraction, we added 1mL of HPLC-grade methanol to each glass vial, vortexed for 5 s, left to stand for 10 min, and vortexed a second time for 5 s. We then stored the glass vials at -20 °C for seven days. Subsequently, we vortexed each vial for 10 s, waited for 10 min, and vortexed again for 10 s. We then removed the swabs using tweezers and squeezed the excess methanol. We dried the samples under nitrogen gas ($N_2$; N-EVAP 111, Organomation, Berlin, MA, USA). We resuspended dried extracts in 100 µL of methanol containing 1 µg of nicotine, used as an internal standard (1.0 mg/mL, Sigma-Aldrich, St Louis, MO, USA, cat no. N5511-1ML), vortexed for 10 s, and transferred them into glass inserts (150 ul, Agilent Technologies, Santa Clara, CA, USA; cat no. 5183-2088). We performed gas chromatography/mass spectrometry (GC-MS) for alkaloid detection following the chromatographic method of Gonzalez et al (2021). Samples (1 uL each) were injected in splitless mode into a Shimadzu GC-MS-QP2020 equipped with an HP-5MS capillary column (30 m × 0.25 mm ID, 0.25 µm; Agilent Technologies, Santa Clara, CA, USA; cat. no. 19091S-433). The injector was held at 250 °C, using helium as the carrier gas at 1mL/min, with a solvent cut time of 4 min. The oven began at 40 °C for 3 min, then ramped to 100 °C at a rate of 6°C/min and held for 1 min. Then, the temperature increased to 200 °C at 4 °C/min and held for 1 min, followed by a final ramp to 300 °C at 20 °C/min with a 5-min hold, yielding a total run time of 50 min. Compounds were ionized by electronic ionization (EI) at 70 eV. The ion source was set to 150 °C, the interface to 230 °C, and the mass spectrometer operated in full scan mode over a mass range from 40 to 500 m/z at a scan rate of 5 scan/s, over the entire GC program. To minimize potential batch effects, we randomized the order of sample analysis so that each batch included individuals from all species and seasons within the same geographical location. As a result, we run *A. trivittata* separately from the other three species. For validation of the analyses, to trace contaminants from the swab materials, 14 field blank runs were performed using the same extraction protocol and chromatographic conditions. Additionally, linear alkanes of the series C8–C20 (Sigma, St Louis, MO, USA) were used for the determination of experimental retention indexes (RI exp).

We exported the GC-MS data files as CDF files and used the Global Natural Products Social Molecular Networking (GNPS) pipelines to perform the deconvolution and library search following the workflow designed by (Aksenov et al. 2021). Converted files were uploaded and shared in the MassIVE online repository (ftp://massive-ftp.ucsd.edu/v11/MSV000100180/). The deconvolution process generated a compound table with molecular features as rows and samples as columns. We selected an intensity of 150,000 as the threshold reliable to differentiate the signal from noise. Data pre-processing was completed by filtering the table through a series of criteria: (1) removing features with a balance score < 60. Balance score ranges from 0–100 and correlates with the quality of the deconvoluted mass spectra; the higher the value, the higher the quality. (2) We also excluded features where the maximum intensity was < 150,000, and (3) removed features associated with swab contaminants (e.g., features where intensity on six blanks was above 150,000). The library search was obtained by comparing our mass spectra with reference libraries of natural products (NIST, Wiley, University of CORSICA) and manual comparison with the Daly et al. (2005) database. For our last filtration step (4), we removed those features where the annotation was a compound different from an alkaloid. Then, we reviewed the automatic GNPS library annotations by manually comparing the experimental mass spectra with those found in the reference libraries. We reported level 2 annotations for each metabolite, which include: experimental retention time (RT), base peak of the mass spectra (m/z), highest peak from the mass spectra (m/z), balance score, cosine similarity, experimental retention index (RI exp), conversion to retention time (RT) to Daly et al. (2005) (Supplementary Table S2). The higher the cosine, the higher the reliability of the putative annotation. Using this information and the characteristic fragments for each alkaloid class (see Gonzalez et al. 2023), each annotated compound was assigned to an alkaloid class following Daly et al. (2005) classification.

The efficiency of the alkaloid collection method varied between species. In the wet season, 22 out of 26 samples (84%) for *A. trivittata* had detected alkaloids, 8 out of 26 samples (31%) for *A. macero* had detected alkaloids, and 7 out of 16 samples (44%) for *A. shihuemoy* had detected alkaloids. In the dry season, 12 out of 13 samples (92%) for *A. trivittata* had detected alkaloids, 3 out of 19 samples (16%) for *A. macero* had detected alkaloids, and 4 out of 6 samples (67%) for *A. shihuemoy* had detected alkaloids.

We used a non-lethal collection method based on the principle of the transcutaneous amphibian stimulator (TAS) (Grant and Land 2002). This approach minimizes the impact on natural populations, specifically for species of conservation concern. Previous studies have shown the effectiveness of this method in other poison frogs (Clark et al. 2006; Hantak et al. 2013; Schulte et al. 2017). Although it does not capture similar abundances to frog skin samples, the quantities obtained are proportional to the total amount present in whole-skin extracts (Basham et al. 2021). Moreover, the effectiveness of the stimulation in inducing alkaloid secretion appears to be species dependent. In our study, 16% of *A. macero* samples collected during the dry season and 31% during the wet season contained detectable alkaloids, whereas the method was more effective for the other two species. This difference may reflect variation in body size, because the largest species, *A. trivittata*, had up to 92% of samples containing alkaloids. However, the smallest species, *A. shihuemoy,* also showed up to 44% of samples with detectable alkaloids, suggesting that body size alone does not fully explain differences in method efficiency. Therefore, although our findings indicate no seasonal differences in overall alkaloid profiles, they should be interpreted with caution due to the potential underestimation of alkaloid quantities and possible species or size-related differences in secretion stimulation efficiency.

**Supplementary Methods S7: Quantification of space use**

To investigate the effect of season on space use area and whether this effect depended on sex, we fitted linear models including the interaction between sex and season for each species. We included snout-vent length (SVL) as a covariate to account for body-size differences that may influence movement capacity, and tracking duration (days tracked) to correct for variation in sampling effort among individuals. Because most individuals were sampled in a single season, seasonal comparisons reflect population-level differences rather than within-individual changes. We evaluated whether the inclusion of individual identity as a random effect was justified. However, the variance associated with individual identity was negligible and models showed singular fits, indicating that the number of recaptures was insufficient to estimate individual-level variance. As a result, individual identity was not included as a random effect in the final models.

Before model fitting, we assessed multicollinearity among predictors using the variance inflation factor *vif()* function from the *car* R package (Fox et al. 2019), and log-transformed space use to meet model assumptions of normality. Model assumptions were evaluated using residual diagnostic plots, and model selection was based on Akaike's Information Criterion (AIC). The significance of main effects and interactions was tested using type II and III Wald $\chi2$ statistics implemented with the *Anova()* function from the *glmmTMB* R package (Brooks et al. 2026). For significant effects, we computed estimated marginal means using the *emmeans ()* and *contrast ()* functions from the *emmeans* R package (Lenth et al. 2025) with the false discovery rate (FDR) correction for multiple testing.

**Supplementary Methods S8: Space use overlap and centroid shifts**

We quantified seasonal shifts in location for individuals recaptured across seasons using two metrics: space use overlap and centroid shifts. Space use overlap was calculated as the ratio between the shared area across seasons and the total area combining both dry and wet seasons, thus providing a measure of the proportion of space shared between seasons. Centroid shifts were calculated as the straight line distance between seasonal space use centroids and used to quantify shifts in the core area of activity. Overlap values of zero indicate complete turnover in space use, whereas values of one indicate identical space use. Lower centroid shift values indicate minor shifts in activity centers, whereas higher values indicate substantial spatial relocation.

**Supplementary Methods S9: Analyses of space use area including tadpole transport movements**

Tadpole transport is an important component of poison frog reproductive behavior and may substantially increase movement distances. Thus, we conducted complementary analyses that included all recorded tadpole transport locations when estimating individual space use (95% kernel density estimates). Tadpole transport observations were unevenly distributed among individuals because these events are difficult to detect consistently during telemetry surveys. Consequently, including these movements may overestimate space use for individuals in which transport events were observed and reduce comparability among individuals and seasons. For this reason, tadpole transport movements were excluded from the analyses presented in the main manuscript.

**Supplementary Methods S10: Daily movement**

We assessed the effect of environmental variables (e.g., daily rainfall and mean daily temperature) on individual daily movement for each species using linear mixed-effect models. We log-transformed daily movement to meet model assumptions of normality. We included daily rainfall, mean daily temperature, sex, and snout-vent length (SVL) as covariates. We also included individual ID nested within season as a random effect to account for repeated measurements of individuals. Before model fitting, we assessed multicollinearity among predictors using the variance inflation factor *vif()* function from the *car* R package. For each species, we compared models including combinations of environmental and individual traits using Akaike's Information Criterion (AIC) to select the best model. We evaluated model assumptions using the *simulateResiduals()* function from the *DHARMa* R package.

**Supplementary Methods S11: Microhabitat use**

We assessed whether the frequency of substrate use (counts) differed between the dry and wet seasons and whether these seasonal differences were influenced by sex, and substrate type (counts ~ season * sex * substrate) for each species. As we have multiple records per individual frog, we included individual ID as a random effect to account for non-independence. To control for differences in sampling effort, we included the logarithm of the total number of observations as an offset term. We compared model performance using Akaike's Information Criterion (AIC), dispersion diagnostics using the *check_overdispersion()* function from the *performance* R package (Lüdecke et al. 2025), and residual plots using the *simulateResiduals()* function in the *DHARMa* R package (Hartig et al. 2024). For *A. femoralis,* the model containing the full season*sex*microhabitat interaction was best supported. For *A. trivittata*, *A.macero*, and *A.shihuemoy*, the best supported model included season*microhabitat interaction, without sex effects. We assessed the significance of main effects and interactions using type III Wald $\chi2$ statistics. We then computed estimated marginal means from the best-fitting models, followed by pairwise contrasts with false discovery rate (FDR) correction for multiple testing. For *A. femoralis,* pairwise comparisons were conducted between seasons within each sex*microhabitat combination, whereas for the other three species, comparisons were made between seasons within each microhabitat category. For each microhabitat, we extracted the ratio of dry to wet seasonal use and its 95% confidence interval. Ratios were derived from back-transformed estimated marginal means. We log-transformed the ratios for visualization, such that positive values indicate greater use during the dry season, and the negative values indicate greater use during the wet season.

### Supplementary Tables

**Table S1. Summary of sample sizes by method (space use, microhabitat use, hormone sampling, and alkaloid profiling).** Values represent samples/individuals per species collected in the dry and wet seasons, with the number of females and males shown in parentheses (females/males).

| Method | Species | Dry Season | Wet Season |
|---|---|---|---|
| Space Use | *A. femoralis* | 26 (12/14) | 30 (7/23) |
| | *A. trivittata* | 16 (7/9) | 20 (8/12) |
| | *A. macero* | 21 (8/13) | 33 (11/22) |
| | *A. shihuemoy* | 7 (6/1) | 17 (8/9) |
| Microhabitat Use | *A. femoralis* | 36 (13/23) | 34 (8/26) |
| | *A. trivittata* | 18 (7/11) | 20 (8/12) |
| | *A. macero* | 35 (11/24) | 35 (12/23) |
| | *A. shihuemoy* | 15 (11/4) | 19 (9/10) |
| Hormones (Testosterone) | *A. femoralis* | 18 (5/13) | 19 (6/13) |
| | *A. trivittata* | 12 (5/7) | 15 (6/9) |
| | *A. macero* | 15 (5/10) | 21 (9/12) |
| | *A. shihuemoy* | 3 (3/0) | 13 (5/8) |
| Hormones (Corticosterone) | *A. femoralis* | 18 (6/12) | 20 (6/14) |

| | | | |
|---|---|---|---|
| | *A. trivittata* | 12 (4/8) | 17 (9/8) |
| | *A. macero* | 17 (5/12) | 24 (10/14) |
| | *A. shihuemoy* | 6 (6/0) | 10 (7/3) |
| Alkaloids | *A. femoralis* | 19 (5/14) | 21 (5/16) |
| | *A. trivittata* | 13 (6/7) | 26 (10/16) |
| | *A. macero* | 19 (7/12) | 26 (9/17) |
| | *A. shihuemoy* | 6 (6/0) | 16 (7/9) |

Values are total n per season, with females/males in parentheses.

Note: Only one male *A. shihuemoy* was recorded during the dry season because males are relatively small compared to the other species, and the tag size was initially too large, leading to frequent frog loss. Improvements in tag size and attachment methods during the wet season increased our number of tracked males.

**Table S2. Level 2 annotations for each metabolite,** including feature ID, alkaloid family, compound name, library match, base peak (m/z), highest peak (m/z), balance score, cosine similarity, Kovats retention index (KI), experimental retention time (RT; min), reported retention times from Daly et al. (2005), conversion to Daly retention time, delta RT, and alkaloid superfamily.

| Feature ID | Family | Compound Name | Compound Name Daly | Library match | Base peak | Highest peak | Balance score | Cosine | Kovats Index | Rt | Rt mins | Daly RT | Daly RT 2 | Daly RT 3 | Conversion to RT Daly | Delta RT | Superfamily |
|---|---|---|---|---|---|---|---|---|---|---|---|---|---|---|---|---|---|
| 619 | 3,5-P | 3,5-Dibutylhexahydro-1H-pyrrolizine | 223B 3,5-Pyrrolizidine | NIST | 166 | 223 | 100 | 0.93 | 1536.05 | 1586.4 | 26.44 | 9.4 | 9.8 | | 9.37 | 0.03 | pyrrolizidine |
| 655 | 3,5-P | (3S,5R,7aS)-3-Pentyl-5-propylhexahydro-1H-pyrrolizine | 223M 3,5-Pyrroilizidine cis | NIST | 152 | 223 | 100 | 0.89 | 1576.05 | 1656 | 27.6 | 8.59 | | | 9.74 | 1.15 | pyrrolizidine |

| | | | | | | | | | | | | | | | | | |
|---|---|---|---|---|---|---|---|---|---|---|---|---|---|---|---|---|---|
| 621 | 5,6,8-I | 223C 5,6,8-Indolizidine | 223C 5,6,8-Indolizidine | Daly | 152 | 223 | 100 | | 1541.22 | 1595.4 | 26.59 | 10.59 | | | 9.42 | 1.17 | indolizidine |
| 944 | 5,6,8-I | (5R,6R,8R,8aS)-6,8-Dimethyl-5-((Z)-non-4-en-8-yn-1-yl)octahydroindolizine | 273A 5,6,8-Indolizidine | NIST | 152 | 273 | 100 | 0.89 | 2045.16 | 2379.6 | 39.66 | 12.86 | 12.95 | | 13.61 | 0.75 | indolizidine |
| 812 | 5,8-I | (5R,8R,8aS)-8-Butyl-5-pentyloctahydroindolizine | 251N 5,8-Indolizidine (5,9Z) | NIST | 180 | 251 | 100 | 0.88 | 1810.74 | 2038.8 | 33.98 | 11.62 | | | 11.79 | 0.17 | indolizidine |

| | | | | | | | | | | | | | | | | |
|---|---|---|---|---|---|---|---|---|---|---|---|---|---|---|---|---|
| 811 | DHQ | (2S,4aS,5R,8aR)-5-(Pent-4-en-1-yl)-2-propyldecahydroquinoline | 249D Decahydroquinoline cis | NIST | 206 | 249 | 100 | 0.89 | 1806.78 | 2032.8 | 33.88 | 11.73 | 11.73 | 12.03 | 11.75 | 0.02 | decahydroquinoline |
| 822 | DHQ | (2S,4aS,5R,8aS)-2-Allyl-5-((Z)-pent-2-en-4-yn-1-yl)decahydroquinoline | 243A Decahydroquinoline cis | NIST | 202 | 243 | 100 | 0.96 | 1832.14 | 2071.2 | 34.52 | 12.32 | 12.68 | 12.74 | 11.96 | 0.36 | decahydroquinoline |
| 838 | DHQ | Isomer of (2R,4aS,5S,8aS)-2-Allyl-5-((Z)-pent-2 | 243A Decahydroquinoline trans | NIST | 202 | 243 | 100 | 0.87 | 1867.42 | 2124.6 | 35.41 | 12.32 | 12.68 | 12.74 | 12.24 | 0.44 | decahydroquinoline |

| | | | | | | | | | | | | | | | | |
|---|---|---|---|---|---|---|---|---|---|---|---|---|---|---|---|---|
| | | -en-4-yn-1-yl)decahydroquinoline | | | | | | | | | | | | | | |
| 845 | DHQ | Isomer of (2R,4aS,5S,8aS)-2-Allyl-5-((Z)-pent-2-en-4-yn-1-yl)decahydroquinoline | 243A Decahydroquinoline 5-epi-trans | NIST | 202 | 243 | 100 | 0.9 | 1875.74 | 2137.2 | 35.62 | 12.32 | 12.68 | 12.74 | 12.31 | 0.43 | decahydroquinoline |
| 982 | DHQ | 267L Decahydroquinoline cis | 267L Decahydroquinoline cis | Daly | 202 | 266 | 100 | | 2143.13 | 2486.4 | 41.44 | 13.97 | 14.5 | | 14.18 | 0.21 | decahydroquinoline |
| 907 | HTX | Histrionicotoxin 259 | 259A Histrionicotoxin | NIST | 96 | 259 | 100 | 0.78 | 1973.26 | 2280 | 38 | 13.2 | | | 13.07 | 0.13 | histrionicotoxin |

| | | | | | | | | | | | | | | | | | |
|---|---|---|---|---|---|---|---|---|---|---|---|---|---|---|---|---|---|
| 921 | HTX | (2R,6R,7S,8S)-7-(But-3-en-1-yl)-2-propyl-1-azaspiro[5.5]undecan-8-ol | 265E Histrionicotoxin | NIST | 152 | 265 | 100 | 0.91 | 2001.54 | 2320.8 | 38.68 | 13.34 | | | 13.29 | 0.05 | histrionicotoxin |
| 939 | HTX | Isomer of Histrionicotoxin 259 | 259A Histrionicotoxin isomer | NIST | 96 | 259 | 100 | 0.88 | 2026.46 | 2354.4 | 39.24 | 13.2 | | | 13.47 | 0.27 | histrionicotoxin |
| 998 | HTX | Octahydrohistrionicotoxin | 291A Histrionicotoxin | Wiley | 178 | 291 | 100 | 0.79 | 2192.75 | 2524.2 | 42.07 | 15.62 | | | 14.38 | 1.24 | histrionicotoxin |
| 999 | HTX | .delta.-17-trans-Histrionicotoxin 283a' | 283A Histrionicotoxin | NIST | 96 | 283 | 69 | 0.77 | 2196.69 | 2527.2 | 42.12 | 15.43 | | | 14.39 | 1.04 | histrionicotoxin |

| | | | | | | | | | | | | | | | | | |
|---|---|---|---|---|---|---|---|---|---|---|---|---|---|---|---|---|---|
| 1003 | HTX | Dihydrohistrionicotoxin 285e | 285E Histrionicotoxin | NIST | 96 | 285 | 100 | 0.78 | 2207.74 | 2533.8 | 42.23 | 15.72 | | | 14.43 | 1.29 | histrionicotoxin |
| 1011 | HTX | Histrionicotoxin 285A | 285A Histrionicotoxin | NIST | 96 | 285 | 100 | 0.79 | 2247.61 | 2554.8 | 42.58 | 15.75 | | | 14.54 | 1.21 | histrionicotoxin |
| 1012 | HTX | Isotetrahydrohistrionicotoxin 287a | 287A Histrionicotoxin | Wiley | 96 | 287 | 70 | 0.67 | 2256.72 | 2559.6 | 42.66 | 16 | | | 14.57 | 1.43 | histrionicotoxin |
| 895 | N-methyl-DHQ | (2R,4aS,5S,8aR)-2-Allyl-1-methyl-5-((Z)-pent-2-en-4-yn-1-yl)decahydroquinoline | 257A N-Methyl-decahydroquinoline cis of 243A | NIST | 216 | 256 | 100 | 0.91 | 1949.23 | 2245.2 | 37.42 | 12.85 | | | 12.89 | 0.04 | N-Methyl-decahydroquinoline |

| | | | | | | | | | | | | | | | | | |
|---|---|---|---|---|---|---|---|---|---|---|---|---|---|---|---|---|---|
| 899 | N-methyl-DHQ | Isomer of (2R,4aS,5S,8aR)-2-Allyl-1-methyl-5-((Z)-pent-2-en-4-yn-1-yl)decahydroquinoline | 257A N-Methyl-decahydroquinoline trans of 243A | NIST | 216 | 256 | 100 | 0.89 | 1957.51 | 2257.2 | 37.62 | 12.85 | | | 12.95 | 0.1 | N-Methyl-decahydroquinoline |
| 1015 | N-methyl-DHQ | (2S,4aS,5R,8aS)-1-Methyl-5-((Z)-pent-2-en-4-yn-1-yl)-2-(penta-3,4-dien-1-yl)decahydroquinoline | 283F N-Methyl-decahydroquinoline of 269AB | NIST | 218 | 284 | 100 | 0.64 | 2264.69 | 2563.8 | 42.73 | | | | 14.59 | ND | N-Methyl-decahydroquinoline |

| | | | | | | | | | | | | | | | | | |
|---|---|---|---|---|---|---|---|---|---|---|---|---|---|---|---|---|---|
| 689 | Tricyclic | 235AA Tricyclic | 235AA Tricyclic | Daly | 192 | 235 | 96 | 0.6 | 1622.96 | 1735.8 | 28.93 | 9.35 | | | 10.17 | 0.82 | Tricyclic |
| 552 | Unclass | 209G - Unclass | 209G Unclass | Daly | 138 | 209 | 79 | | 1447.76 | 1428.6 | 23.81 | 7.36 | 7.93 | | 8.53 | 0.6 | Unclass |
| 788 | Unclass | Unclass | Unclass | Daly | 194 | 251 | 100 | | 1765.67 | 1968 | 32.8 | | | | 11.41 | NA | Unclass |
| 886 | Unknown | Unknown | Unknown | Daly | 166 | 267 | 100 | | 1932.65 | 2221.2 | 37.02 | | | | 12.76 | NA | Unknown |
| 709 | Pyr | 225C 2,5-Pyrrolidine | 225C 2,5-Pyrrolidine | Daly | 126 | 254 | 100 | | 1648.27 | 1777.8 | 29.63 | 9.44 | | | 10.39 | 0.95 | Pyr |

**Table S3. Predictors of space use (HPI95)** Linear models testing effects of season, sex, body size (SVL), and tracking duration on log-transformed 95% space use. Significant predictors (p < 0.05) are shown in bold.

| Species | Predictor | β | SE | t | p | |
|---|---|---|---|---|---|---|
| *Allobates femoralis* | Season (wet) | 0.111 | 0.159 | 0.700 | **0.014** | * |
| | Sex (male) | -0.199 | 0.149 | -1.340 | 0.111 | |
| | SVL | 0.159 | 0.091 | 1.740 | 0.087 | |
| | Days tracked | 0.097 | 0.037 | 2.590 | **0.012** | * |
| | Season × Sex | 0.529 | 0.147 | 3.600 | **<0.001** | *** |
| *Ameerega trivittata* | Season (wet) | -0.374 | 0.149 | -2.510 | **0.017** | * |
| | Sex (male) | 0.222 | 0.150 | 1.480 | 0.149 | |
| *Ameerega macero* | Season (wet) | -0.105 | 0.130 | -0.810 | 0.069 | |
| | Sex (male) | -0.694 | 0.197 | -3.520 | **<0.001** | *** |
| | SVL | 0.273 | 0.094 | 2.920 | **0.005** | ** |
| | Season × Sex | -0.283 | 0.131 | -2.170 | **0.035** | * |
| *Ameerega shihuemoy* | Season (wet) | 0.369 | 0.237 | 1.560 | 0.134 | |
| | SVL | -0.229 | 0.129 | -1.770 | 0.091 | |

**Significance codes:**
*** $p < 0.001$, ** $p < 0.01$, * $p < 0.05$, $p < 0.1$

**Table S4. Pairwise contrasts of seasonal differences in space use(log-transformed)** estimated as dry–wet season differences within each species × sex group, obtained from the estimated marginal means (emmeans). Positive estimates indicate larger space use in the dry season, and negative estimates indicate smaller space use in the dry season. Significant contrasts (p < 0.05) are shown in bold.

| Species | Sex | Contrast | Estimate | SE | t-value | p-value | |
|---|---|---|---|---|---|---|---|
| *Allobates femoralis* | f | Dry − Wet | 1.278 | 0.501 | 2.550 | **0.014** | * |
| | m | Dry − Wet | -0.836 | 0.350 | -2.390 | **0.021** | * |

| | | | | | | | |
|---|---|---|---|---|---|---|---|
| *Ameerega trivittata* | all | Dry − Wet | -0.748 | 0.298 | -2.510 | **0.017** | * |
| *Ameerega macero* | f | Dry − Wet | -0.778 | 0.418 | -1.860 | 0.069 | |
| | m | Dry − Wet | 0.356 | 0.310 | 1.150 | 0.256 | |
| *Ameerega shihuemoy* | — | No significant seasonal contrast | — | — | — | — | |

**Significance codes:**
*** $p < 0.001$, ** $p < 0.01$, * $p < 0.05$, $p < 0.1$

**Supplementary Results S1: Space use overlap and centroid shifts**

Seasonal space use overlap and centroid shifts varied among species and sexes (Supplementary Tables S5 and S6). In the seasonal breeder *A. femoralis*, females showed near-zero overlap between seasons (0.1% and 0.08%; n=2), whereas males showed low overlap (19% and 21%; n=2), with relatively small centroid shifts (< 7 m in both sexes). In contrast, *A. trivittata* (n = 1 female, 1 male) showed complete seasonal turnover in space use in both individuals (overlap = 0%), accompanied by large centroid shifts (23.51 m in the male, and 100.49 in the female). *A. macero* exhibited similar low levels of overlap (females: n = 3, mean ± SD = 11% ± 4%; males: n = 4, 11% ± 8%), although centroid shifts were more variable, particularly among males (females: 2.44 ± 0.81 m; males: 8.92 ± 13.74 m). The single recaptured individual of *A. shihuemoy* showed low overlap (7%) and a small centroid shift (2.36 m). These results should be interpreted cautiously due to the limited sample sizes.

**Table S5 Individual-level space use overlap and centroid shifts between recaptures for each species and sex.** Overlap represents the proportional overlap in space use between individuals, and centroid shifts (m) is the straight line distance between the individuals' centers of activity.

| Species | Sex | Recapture ID | Space Use Overlap | Centroid shifts (m) |
|---|---|---|---|---|
| *Allobates femoralis* | female | af_f01 | 0.001 | 6.981 |
| | | af_f02 | 0.001 | 3.147 |
| | male | af_m01 | 0.187 | 3.001 |
| | | af_m02 | 0.209 | 3.273 |
| *Ameerega trivittata* | female | at_f01 | 0.000 | 100.487 |
| | male | at_m01 | 0.000 | 23.516 |
| *Ameerega macero* | female | am_f01 | 0.098 | 1.827 |
| | | am_f02 | 0.148 | 3.356 |
| | | am_f03 | 0.069 | 2.136 |
| | male | am_m01 | 0.200 | 1.570 |
| | | am_m02 | 0.111 | 4.134 |
| | | am_m03 | 0.000 | 29.401 |
| | | am_m04 | 0.122 | 0.558 |
| *Ameerega shihuemoy* | female | as_f01 | 0.072 | 2.365 |

**Table S6. Mean ± SD of space use overlap and centroid shifts (m) calculated for each species and sex.**

| Species | Sex | n | Space Use Overlap mean ± SD | Centroid shifts (m) mean ± SD |
|---|---|---|---|---|
| *Allobates femoralis* | female | 2 | 0.001± 0.000 | 5.064 ± 2.711 |

| | | | | |
|---|---|---|---|---|
| | male | 2 | 0.198 ± 0.015 | 3.137 ± 0.193 |
| *Ameerega trivittata* | female | 1 | 0.000 | 100.487 |
| | male | 1 | 0.000 | 23.516 |
| *Ameerega macero* | female | 3 | 0.105 ± 0.040 | 2.440 ± 0.808 |
| | male | 4 | 0.108 ± 0.082 | 8.916 ± 13.740 |
| *Ameerega shihuemoy* | female | 1 | 0.072 | 2.365 |

**Supplementary Results S2: Space use area including tadpole transport movements**

The inclusion of tadpole transport movements in the space use area models altered some statistical relationships but did not qualitatively change the main biological interpretations for most species. In *A. femoralis*, the previously significant seasonal expansion in male space use area during the wet season was no longer statistically significant, although the directional trend remained similar. In *A. trivittata* and *A. macero*, inclusion of tadpole transport movements did not qualitatively change the results. In *A. shihuemoy*, seasonal differences in space use area became significant when tadpole transport locations were included.

**Table S7** Post hoc seasonal contrasts in space use after including tadpole transport movements in kernel density estimates (95% utilization distributions). Positive estimates indicate greater dry-season space use, whereas negative estimates indicate greater wet-season space use.

| Species | Sex | Dry − Wet estimate | SE | 95% CI | p-value | Interpretation |
|---|---|---|---|---|---|---|
| *Allobates femoralis* | All | 0.540 | 0.340 | -0.142, 1.222 | 0.118 | No significant seasonal difference |

| | | | | | | |
|---|---|---|---|---|---|---|
| | Female | 1.661 | 0.539 | 0.579, 2.743 | **0.003** | Larger dry-season space use |
| | Male | -0.581 | 0.376 | -1.337, 0.175 | 0.129 | Wet-season expansion no longer significant |
| *Ameerega trivittata* | All | -0.906 | 0.367 | -1.653, -0.159 | **0.019** | Larger wet-season space use |
| | Female | 0.206 | 0.521 | -0.859, 1.271 | 0.696 | No significant seasonal difference |
| | Male | -0.683 | 0.438 | -1.577, 0.211 | 0.129 | No significant seasonal difference |
| *Ameerega macero* | All | -0.198 | 0.260 | -0.719, 0.324 | 0.449 | No significant seasonal difference |
| | Female | -0.777 | 0.419 | -1.618, 0.065 | 0.070 | Marginal wet-season expansion |
| | Male | 0.381 | 0.310 | -0.242, 1.004 | 0.225 | No significant seasonal difference |
| *Ameerega shihuemoy* | All | 1.631 | 0.480 | 0.631, 2.630 | **0.003** | Larger dry-season space use |
| | Female | 0.996 | 0.591 | -0.233, 2.224 | 0.107 | No significant seasonal difference |

| | | | | | | |
|---|---|---|---|---|---|---|
| | Male | 0.454 | 1.203 | -2.049, 2.956 | 0.710 | No significant seasonal difference |

**Table S8. Environmental predictors of daily movement distance.** Linear mixed models testing effects of rainfall and temperature on log-transformed daily movement distance. Individual-season identity (ID_season) was included as a random effect. Wald $\chi^2$ statistics and associated p-values were obtained from Type II analysis of deviance tests.

| Species | Predictor | Estimate (β) | SE | Wald ($\chi^2$) | p-value | |
|---|---|---|---|---|---|---|
| *Allobates femoralis* | Daily rainfall | 0.006 | 0.002 | 9.980 | **0.002** | ** |
| | Mean daily temperature | 0.001 | 0.030 | 0.002 | 0.967 | |
| *Ameerega trivittata* | Daily rainfall | 0.002 | 0.002 | 0.420 | 0.516 | |
| | Mean daily temperature | -0.084 | 0.037 | 5.310 | **0.021** | * |
| *Ameerega macero* | Daily rainfall | 0.001 | 0.002 | 0.631 | 0.427 | |
| | Mean daily temperature | 0.068 | 0.025 | 7.500 | **0.006** | ** |
| | Sex (male) | 0.491 | 0.159 | 9.550 | **0.002** | ** |
| | Body size (SVL) | 0.091 | 0.037 | 5.952 | **0.015** | * |
| *Ameerega shihuemoy* | Daily rainfall | 0.003 | 0.003 | 1.460 | 0.227 | |
| | Mean daily temperature | 0.072 | 0.052 | 1.900 | 0.168 | |

**Significance codes:**
*** $p < 0.001$, ** $p < 0.01$, * $p < 0.05$, $p < 0.1$

**Table S9. Type III Wald $\chi^2$ tests from generalized linear mixed models for microhabitat (negative binomial, log offset).** Models assessing the main effects and interactions of season, sex, and substrate category on substrate-use counts for the seasonal breeders (*Allobates femoralis* and *Ameerega trivittata*) and the year-round breeders (*Ameerega macero* and *Ameerega shihuemoy*) poison frogs.

| Species | Effect | $\chi^2$ | df | p-value |
|---|---|---|---|---|

| | | | | | |
|---|---|---|---|---|---|
| ***Allobates femoralis*** | (Intercept) | 26.295 | 1 | **0.000** | *** |
| | season | 2.710 | 1 | 0.100 | . |
| | sex | 1.044 | 1 | 0.307 | |
| | substrate categories | 92.524 | 8 | **< 0.001** | *** |
| | season*sex | 2.464 | 1 | 0.116 | |
| | season*substrate categories | 14.614 | 8 | 0.067 | . |
| | sex*substrate categories | 3.288 | 8 | 0.915 | |
| | season*sex*substrate categories | 22.372 | 8 | **0.004** | ** |
| ***Ameerega trivittata*** | (Intercept) | 14.000 | 1 | **0.000** | *** |
| | season | 0.068 | 1 | 0.795 | |
| | sex | 0.090 | 1 | 0.764 | |
| | substrate categories | 33.329 | 8 | **< 0.001** | *** |
| | season*sex | 0.003 | 1 | 0.959 | |
| | season*substrate categories | 31.921 | 8 | **< 0.001** | *** |
| | sex*substrate categories | 9.208 | 8 | 0.325 | |
| | season*sex*substrate categories | 3.440 | 8 | 0.904 | |
| ***Ameerega macero*** | (Intercept) | 61.241 | 1 | **< 0.001** | *** |
| | season | 0.002 | 1 | 0.963 | |
| | sex | 2.998 | 1 | 0.083 | . |
| | substrate categories | 69.511 | 9 | **< 0.001** | *** |
| | season*sex | 1.034 | 1 | 0.309 | |
| | season*substrate categories | 19.101 | 9 | **0.024** | * |
| | sex*substrate categories | 17.636 | 9 | **0.040** | * |

| | | | | | |
|---|---|---|---|---|---|
| | season*sex*substrate categories | 16.070 | 9 | 0.065 | . |
| ***Ameerega shihuemoy*** | (Intercept) | 51.349 | 1 | **0.000** | *** |
| | season | 0.102 | 1 | 0.749 | |
| | sex | 0.540 | 1 | 0.462 | |
| | substrate categories | 73.244 | 7 | **0.000** | *** |
| | season*sex | 1.227 | 1 | 0.268 | |
| | season*substrate categories | 6.352 | 7 | 0.499 | |
| | sex*substrate categories | 2.125 | 7 | 0.953 | |
| | season*sex*substrate categories | 8.216 | 7 | 0.314 | |

**Significance codes:**
*** $p < 0.001$, ** $p < 0.01$, * $p < 0.05$, . $p < 0.1$

**Table S10. Post-Hoc pairwise comparison of substrate use between dry and wet seasons for species with significant interactions**. Tests were performed on the log scale, and ratios are back-transformed. Significant differences indicate the substrates that were used more or less in one season than the other. Sex was only relevant for *A. femoralis*.

| Species | Substrate | Sex | Contrast | Ratio | SE | z | p-value | |
|---|---|---|---|---|---|---|---|---|
| ***Allobates femoralis*** | On log | female | dry/wet | 0.557 | 0.153 | -2.127 | **0.0334** | * |
| | Treefall refuge | male | dry/wet | 0.185 | 0.151 | -2.074 | **0.0381** | * |
| | Under leaf litter | male | dry/wet | 1.374 | 0.208 | 2.097 | **0.0360** | * |
| | Under log | male | dry/wet | 1.789 | 0.51 | 2.042 | **0.0412** | * |
| ***Ameerega trivittata*** | Inside roots | — | dry/wet | 1.584 | 0.343 | 2.125 | **0.0336** | * |

| | | | | | | | | |
|---|---|---|---|---|---|---|---|---|
| | Potential clutch site | — | dry/wet | 0.529 | 0.124 | -2.710 | **0.0062** | ** |
| | On leaf litter | — | dry/wet | 0.325 | 0.068 | -5.401 | **<0.001** | *** |
| | Treefall refuge | — | dry/wet | 7.399 | 4.548 | 3.256 | **0.0001** | *** |
| | Under leaf litter | — | dry/wet | 2.007 | 0.353 | 3.960 | **<0.001** | *** |
| ***Ameerega macero*** | On log | — | dry/wet | 1.744 | 0.487 | 1.996 | **0.0460** | * |
| | Rock refuge | — | dry/wet | 4.232 | 2.889 | 2.113 | **0.0346** | * |
| | Under log | — | dry/wet | 2.269 | 0.493 | 3.771 | **0.0002** | ** |

**Significance codes:**
*** $p < 0.001$, ** $p < 0.01$, * $p < 0.05$, $p < 0.1$

**Table S11. Species-specific ANOVA/GLM results for testosterone release rates.** Linear models (LM, log-transformed) were used for all species except *Ameerega macero*, which was analyzed with a generalized linear model (GLM) with a Gamma error distribution and log link. F values are shown for LMs, and LR Chisq for the GLM.

| Species | Model | Predictor | Statistic | p-value | |
|---|---|---|---|---|---|
| *Allobates femoralis* | LM (log) | Intercept | 5.754 | 0.022 | . |
| | | season | 14.596 | **0.001** | ** |
| | | sex | 8.262 | **0.007** | ** |
| | | SVL | 0.301 | 0.587 | |
| | | season*sex | 10.947 | **0.002** | ** |
| *Ameerega trivittata* | LM (log) | Intercept | 8.301 | **0.008** | ** |
| | | season | 1.755 | 0.199 | |
| | | sex | 0.208 | 0.653 | |
| | | SVL | 2.617 | 0.120 | |
| | | season*sex | 0.989 | 0.331 | |

| | | | | | |
|---|---|---|---|---|---|
| *Ameerega macero* | GLM Gamma | sex | 0.000 | 0.997 | |
| | | season | 0.144 | 0.704 | |
| | | SVL | 0.534 | 0.465 | |
| | | season*sex | 1.640 | 0.203 | |
| *Ameerega shihuemoy* | LM (log) | season | 2.220 | 0.160 | |
| | | SVL | 5.043 | **0.043** | * |

**Significance codes:**
*** $p < 0.001$, ** $p < 0.01$, * $p < 0.05$, $p < 0.1$

**Table S12. Predictors of space use (HPI95) from testosterone release rates.** Linear models testing effects of testosterone and sex on log-transformed HPI95 across species.

| Species | n | Predictor | Estimate | SE | p-value |
|---|---|---|---|---|---|
| *Allobates femoralis* | 30 | Testosterone | 0.004 | 0.005 | 0.365 |
| | | Sex (male) | -0.105 | 0.979 | 0.915 |
| | | Testosterone * Sex | 0.011 | 0.014 | 0.430 |
| *Ameerega trivittata* | 27 | Testosterone | -0.011 | 0.006 | 0.074 |
| | | Sex (male) | -0.673 | 0.511 | 0.201 |
| | | Testosterone * Sex | 0.011 | 0.006 | 0.108 |
| *Ameerega macero* | 29 | Testosterone | -0.006 | 0.024 | 0.797 |
| | | Sex (male) | 0.293 | 1.148 | 0.800 |
| | | Testosterone * Sex | 0.002 | 0.024 | 0.941 |
| *Ameerega shihuemoy* | 16 | Testosterone | -0.002 | 0.004 | 0.565 |
| | | Sex (male) | 0.589 | 0.523 | 0.281 |

**Table S13. Species-specific ANOVA/GLM results for corticosterone release rates.** Linear models (LM, log-transformed) were used for all species except *Ameerega trivittata*, which was analyzed with a generalized linear model (GLM) with a Gamma error distribution and log link. F values are shown for LMs, and LR Chisq for the GLM.

| **Species** | **Model** | **Predictor** | **Statistic** | **p-value** | |
|---|---|---|---|---|---|
| *Allobates femoralis* | LM (log) | Intercept | 8.404 | **0.006** | ** |
| | | season | 3.143 | **0.085** | * |
| | | sex | 1.208 | 0.279 | |
| | | SVL | 0.001 | 0.982 | |
| | | season*sex | 0.797 | 0.378 | |
| *Ameerega trivittata* | GLM Gamma | sex | 0.141 | 0.707 | |
| | | season | 0.970 | 0.325 | . |
| | | SVL | 1.597 | 0.206 | |
| | | sex*season | 0.123 | 0.726 | |
| *Ameerega macero* | LM (log) | Intercept | 10.580 | **0.002** | ** |
| | | season | 3.782 | 0.059 | . |
| | | sex | 1.021 | 0.319 | |
| | | SVL | 0.076 | 0.785 | |
| | | season*sex | 2.055 | 0.160 | |
| *Ameerega shihuemoy* | LM (log) | season | 0.490 | 0.497 | |
| | | SVL | 1.960 | 0.185 | |

**Significance codes:**
*** $p < 0.001$, ** $p < 0.01$, * $p < 0.05$, $p < 0.1$

**Table S14.** Species-specific PERMANOVA testing for seasonal differences in alkaloid composition using Bray–Curtis dissimilarities of alkaloid family abundances (ng). Reported values include degrees of freedom (Df), pseudo-F statistic (F), proportion of variance explained by season ($R^2$), and permutation-based p-values (999 permutations). No significant seasonal differences were detected within species.

| Species | Df | F | $R^2$ | P-value |
|---|---|---|---|---|
| *Ameerega trivittata* | 1 | 1.404 | 0.042 | 0.212 |

| | | | | |
|---|---|---|---|---|
| *Ameerega macero* | 1 | 1.134 | 0.112 | 0.361 |
| *Ameerega shihuemoy* | 1 | 1.097 | 0.109 | 0.413 |

**Table S15. Seasonal comparisons of alkaloid family abundance within each species.** Differences between dry and wet seasons were assessed using pairwise Wilcoxon rank-sum tests. Reported values include sample sizes (n), test statistic (W), raw and Benjamini–Hochberg-adjusted p-values.

| Species | Alkaloid family | Dry (n) | Wet (n) | W | P-value | P adjusted |
|---|---|---|---|---|---|---|
| *Ameerega trivittata* | 5,8-I | 12 | 22 | 162 | 0.514 | 0.734 |
| | DHQ | | | 212 | 0.204 | 0.478 |
| | HTX | | | 217 | 0.157 | 0.478 |
| | N-methyl-DHQ | | | 211 | 0.214 | 0.478 |
| | Pyr | | | 176 | 0.612 | 0.765 |
| | Tricyclic | | | 196 | 0.263 | 0.478 |
| | Unclass | | | 162 | 0.514 | 0.734 |
| | Unknown | | | 176 | 0.612 | 0.765 |
| *Ameerega macero* | 3,5-P | 3 | 8 | 273 | 0.101 | 0.478 |
| | 5,6,8-I | | | 246 | 0.966 | 1.000 |
| | DHQ | | | 242 | 0.811 | 0.927 |
| | HTX | | | 215 | 0.221 | 0.478 |
| | N-methyl-DHQ | | | 260 | 0.261 | 0.478 |
| | Tricyclic | | | 260 | 0.261 | 0.478 |

| | | | | | | |
|---|---|---|---|---|---|---|
| | Unclass | | | 252 | 0.834 | 0.927 |
| *Ameerega shihuemoy* | 3,5-P | 4 | 7 | 72 | 0.004 | 0.074 |
| | 5,6,8-I | | | 64 | 0.022 | 0.220 |
| | DHQ | | | 58 | 0.298 | 0.497 |
| | HTX | | | 63 | 0.196 | 0.478 |
| | Unclass | | | 48.5 | 1.000 | 1.000 |

**Supplementary figures**

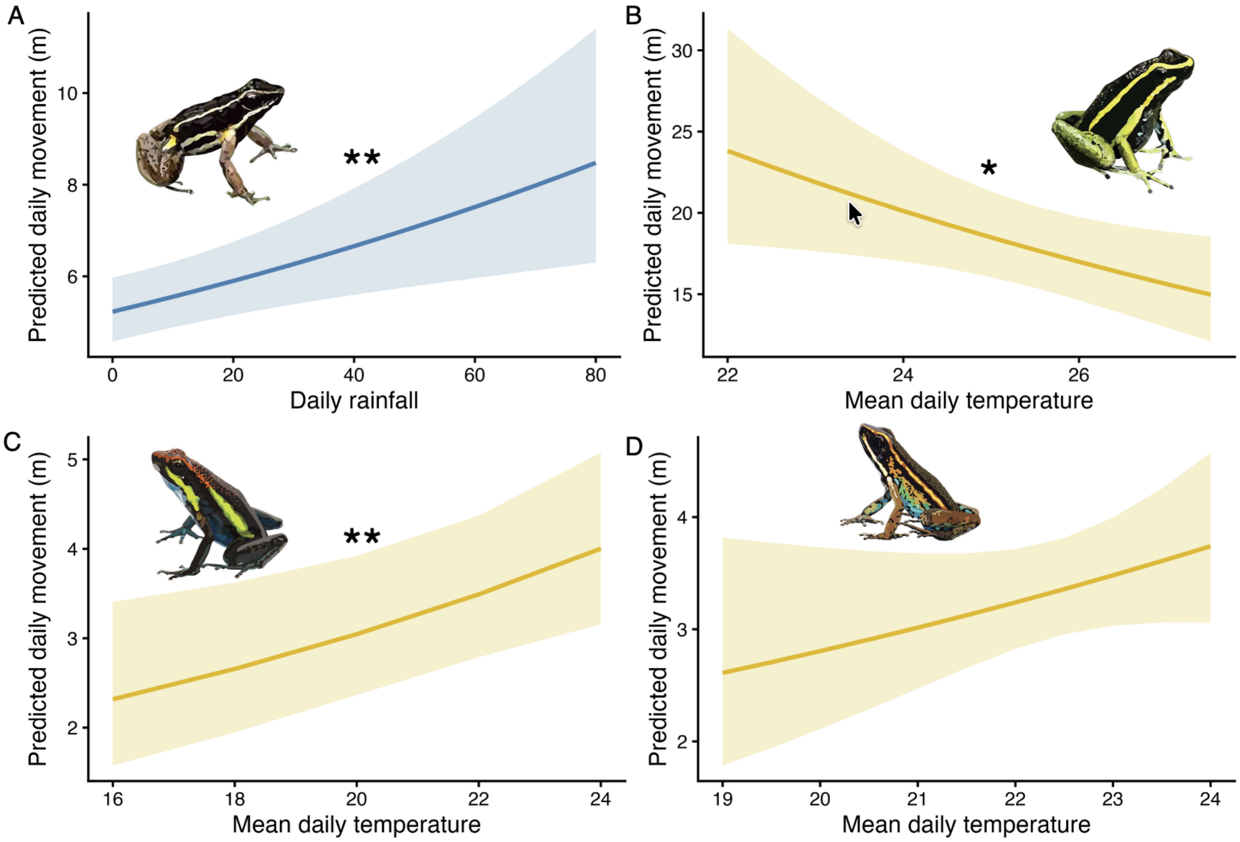


**Fig. S3 Predicted effect of rainfall and temperature on daily movement across poison frog species.** Lines show model-predicted values of log-transformed daily movement distance, and shaded areas represent 95% confidence intervals. (A) In the seasonal breeder *A. femoralis*, daily movement increased with increasing rainfall. (B) In the seasonal breeder *A. trivittata*, daily movement decreased with increasing mean daily temperature. (C) In the year-round breeder *A. macero*, daily movement increased with increasing mean daily temperature. (D) In the year-round breeder *A. shihuemoy*, daily movement also increased with temperature, although this effect was not statistically significant. Blue lines represent rainfall effects and yellow lines represent temperature effects. Significant codes: '**' < 0.01 '*'< 0.05 ''> 0.05

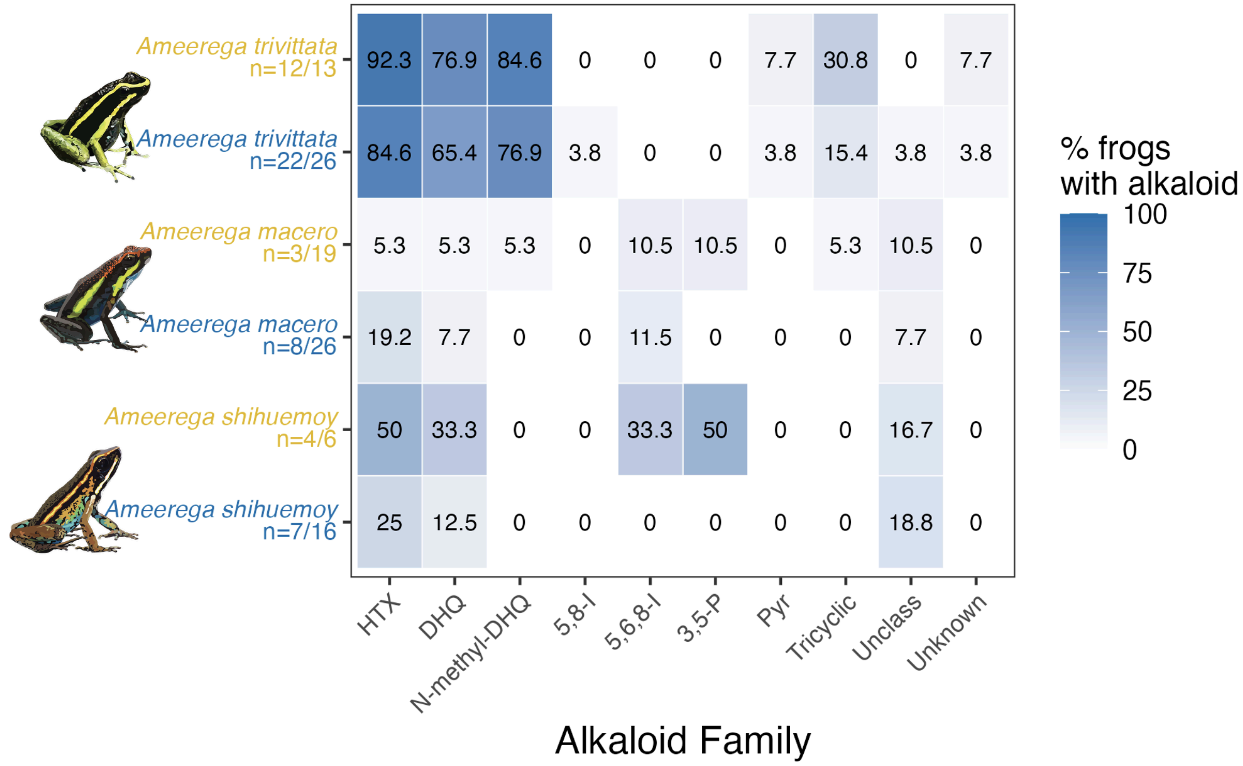


**Fig. S4. Heatmap showing the prevalence (%) of alkaloid families** detected in swabs of *Ameerega trivittata*, *Ameerega macero*, and *Ameerega shihuemoy* by season. Values within cells indicate the percentage of frogs containing at least one alkaloid from each family (sample sizes indicated as samples with detectable alkaloids/total sampled). Darker blue shading indicates higher prevalence. Alkaloid families include histrionicotoxins (HTX), decahydroquinolines (DHQ), N-methyl decahydroquinolines (N-methyl-DHQ), 5,8-disubstituted indolizidines (5,8-I), 5,6,8-trisubstituted indolizidines (5,6,8-I), 3,5-disubstituted pyrrolizidines (3,5-P), piperidines (Pyr), tricyclic alkaloids, unclassified alkaloids, and unknown alkaloids. No alkaloids were detected in the seasonal breeder *Allobates femoralis*.
